# A cyanobacterial screening platform for Rubisco mutant variants

**DOI:** 10.1101/2025.01.22.633911

**Authors:** Ute A. Hoffmann, Anna Z. Schuppe, Axel Knave, Emil Sporre, Hjalmar Brismar, Elias Englund, Per-Olof Syrén, Elton P. Hudson

## Abstract

Rubisco is the main entry point of inorganic carbon into the biosphere and a central player in the global carbon system. Its relatively low catalytic constant as well as its tendency to also accept O_2_ as a substrate have made it a common target of enzyme engineering. We have developed an enzyme engineering and screening platform for Rubisco using the model cyanobacterium *Synechocystis* sp. PCC 6803. Starting with the Form II Rubisco from *Gallionella,* we first show that the enzyme can replace the native Form I Rubisco in *Synechocystis* and that growth rates become sensitive to CO_2_ and O_2_ levels. We address the challenge of designing a zero-shot input library, without prior experimental knowledge, by coupling the phylogenetically-guided model EVmutation with “*in silico* evolution”. Starting with this targeted mutagenesis library, we used competitive growth coupled to deep sequencing to compare the properties of Rubisco protein variants under different cultivation conditions. We identified an amino acid exchange which increased the thermostability of *Gallionella* Rubisco and conveyed resilience to detrimental exchanges. The establishment of this platform is a first step towards high-throughput screening of Rubisco variants in *Synechocystis* and creating optimized enzyme variants to accelerate the Calvin-Benson-Bassham cycle in cyanobacteria and possibly chloroplasts.

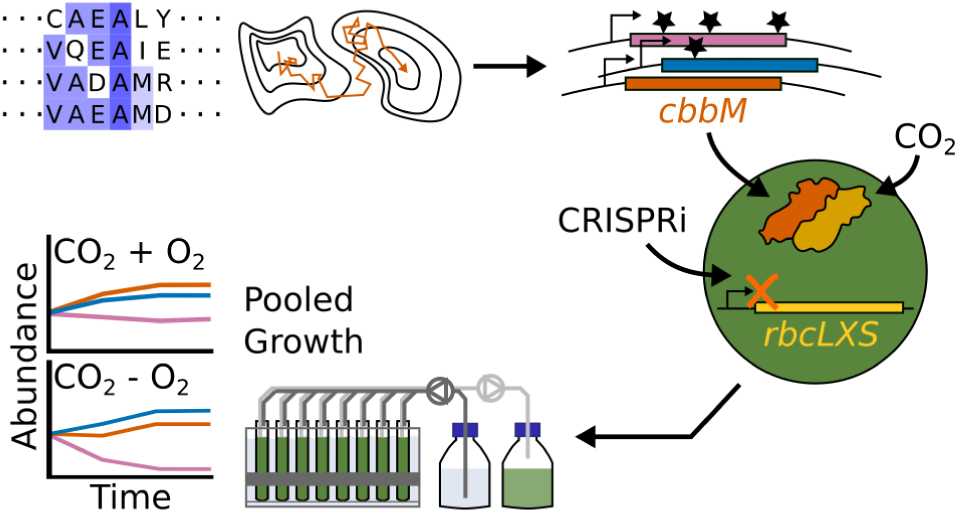

## Introduction

Most inorganic carbon enters the biosphere via fixation by Rubisco as part of the Calvin-Benson-Bassham (CBB) cycle (1, 2). Rubisco has been an important, though elusive target for enzyme engineering in efforts to improve growth, photosynthetic activity and productivity of photoautotrophs (1–6). Despite its central role in carbon metabolism, Rubisco exhibits a low turnover number (*k_cat,C_*) for a central metabolic enzyme (7). Further, in addition to the carboxylation of ribulose-1,5-bisphosphate (RuBP), which results in two molecules of 3-phosphoglycerate (3PG), Rubisco catalyzes the oxygenation of RuBP, leading to one molecule 3PG and the byproduct 2-phosphoglycolate (2PG), which is converted into 3PG by an energy-intensive salvage pathway (photorespiration (1)). Interestingly, there may exist trade-offs between Rubisco’s enzyme parameters (1, 2). For instance, a higher specificity for CO_2_ compared to O_2_ (higher S(℅)) seems to come at the expense of a lower carboxylation turnover number (lower *k_cat,C_*) (1). The specificity factor S(℅) is defined as the ratio of Rubisco’s specificity constants for CO_2_ and O_2_, *k_C_* and *k_O_*, which equals the ratio of the catalytic efficiencies for both substrates, 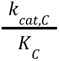 and 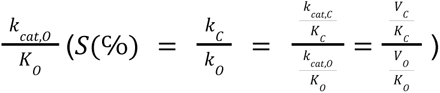. At given concentrations of CO_2_ and O_2_, [CO_2_] and [O_2_], it gives the ratio of the velocities of carboxylation and oxygenation reaction 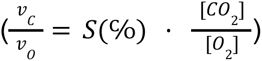. Note that the limiting rate 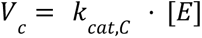 is a theoretical upper boundary of the reaction velocity at a given enzyme concentration [*E*] reached in the absence of substrate limitation and the Michaelis constant for CO^2^, 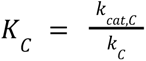, gives a measure for the enzyme’s affinity to its substrate.

Rubisco homologues are classified into Form I, II and III enzymes (1). The basic functional unit of all Rubiscos is a homodimer of two large subunits (L_2_), which may assemble into higher-order oligomers such as tetramers (L_4_) or hexamers (L_6_) (1). Cyanobacterial and plant Rubiscos belong to Form I Rubiscos, which acquired a small subunit through evolution (1, 8). They form heterohexadecimers consisting of eight large and eight small subunits (L_8_S_8_). The small subunit seems to have increased Rubisco selectivity for CO_2_ (1, 8). The need to maintain protein-protein interactions, e.g. with Rubisco-specific assembly factors or the small subunit, might constrain the evolvability of Form I Rubisco large subunits compared to other forms (6, 8, 9).

Compared to Form I Rubiscos, Form II Rubiscos are usually more easily expressed heterologously due to their lower dependence on chaperones (2, 10). The fastest Rubisco reported so far, CbbM, is a Form II Rubisco which forms a dimer (L_2_), and was identified from a *Gallionella* species which was part of a metagenomic assembly (10). Its *k_cat,C_* was measured as 22.2 ± 1.1 s^−1^, which is approximately 6 times as fast as median plant Rubiscos when measured at the same reaction temperature of 25°C (10). However, the increased catalytic constant of *Gallionella* Rubisco comes at the expense of a specificity; S(℅) of 10.0 ± 0.1 was reported, significantly lower than Form I Rubiscos from plants (median 94 ± 11) and the cyanobacterium *Synechococcus elongatus* PCC 6301 (42.7 ± 2.8). In concordance with these low specificity values of Form II Rubiscos, exchanging plant or cyanobacterial Rubisco for Form II Rubisco leads to a high-carbon-dependent phenotype (11, 12).

Attempts to optimize the enzymatic properties of Rubisco have been undertaken for several decades (1, 2). Even though increasing plant growth was often the ultimate goal of these attempts, long generation times, separation of Rubisco-encoding genes *rbcL* and *rbcS* into chloroplast and nuclear genomes, and the difficulties of chloroplast transformations precluded plants from being used as a platform for high-throughput enzyme engineering (1, 2). As a result, several bacterial screening platforms were developed. These included *Escherichia coli* (*E. coli*) strains which were engineered to be dependent on Rubisco activity, and facultative autotrophs such as *Rhodobacter capsulatus* or *Cupriavidus necator* (1). However, screening efforts have had so far only limited success. Although improved expression of tested Rubiscos was achieved, catalytic parameters were rarely improved (1). Cyanobacteria are the only known bacteria performing oxygenic photosynthesis and are evolutionarily closely related to chloroplasts. The model organism *Synechocystis* sp. PCC 6803 (*Synechocystis*) is a unicellular cyanobacterium capable of both photoautotrophic, mixotrophic as well as chemoheterotrophic growth. We considered the strong dependence of metabolite levels on *Synechocystis* growth condition (13–16) an interesting set of constraints for developing new Rubiscos. Hence, we decided to establish a *Synechocystis* host strain to be used in the high-throughput screening of Rubisco variants. We envisioned that, depending on gas feed conditions, this system should enable screening for different enzyme parameters. Compared to *E. coli*, Rubisco is essential in *Synechocystis* and well integrated into its metabolism.

Previous attempts to optimize Rubisco in high-throughput used random input or deep mutational scans (DMS) to create input libraries (1, 2, 17). In a DMS performed for the complete Form II *Rhodospirillum rubrum* (*R. rubrum*) Rubisco, single amino acid exchanges did not lead to a substantially more active enzyme, implying that several amino acid exchanges at once are needed to reach this goal (17). For various other enzymes and proteins, the creation of mutant variants with several amino acid exchanges by introducing these via error-prone PCR usually lead to a strong fitness decrease (18–22). A computationally based “zero-shot prediction”, i.e. a targeted library design, can increase chances of acquiring a functional protein or enzyme and to overcome this problem (23). According to benchmarking on existing substitution data sets, EVmutation (24, 25) performs comparatively well in predicting the fitness of higher-order mutant variants when lacking experimental data (22, 23, 26). EVmutation uses multiple sequence alignments of homologues of a protein of interest to capture both the conservation of a residue at a given position as well as the pair-wise co-variation of amino acid positions (24, 25). To address the challenge of picking residue positions to be included in a higher-order zero-shot library, we coupled predictions by EVmutation with “*in silico* evolution”, a variant of the Metropolis-Hastings algorithm (27). Similar methods were previously successfully employed to generate functional proteins with a high number of amino acid exchanges (28–30).

Previously, we established a growth-competition-based screening system of *Synechocystis*, which we used to interrogate the fitness effect of knocking down every annotated gene using CRISPR interference (CRISPRi) (31, 32). Here, we developed this set-up to apply it to enzyme engineering. We engineered a *Synechocystis* host strain so that its growth became dependent on heterologously expressed Rubisco CbbM. By coupling EVmutation (24, 25) with *in silico* evolution (27), we identified four amino acid exchanges expected to improve CbbM fitness which we combined to create 16 combinatorial higher-order mutant variants. We performed a pooled growth competition of the *Synechocystis* host strain expressing these different variants in different gas feeds. None of the predicted exchanges improved growth of *Synechocystis* compared to CbbM. Two of the tested amino acid exchanges were able to rescue the detrimental effect of the other two investigated replacements, enabling us to map epistatic relationships between different residues. The developed system will enable further screening efforts and to investigate the evolvability of CbbM towards more beneficial enzyme parameters.

## Results and Discussion

### Efficient repression of *Synechocystis* Rubisco, RbcLS, using CRISPR interference

The aim of this study was to engineer a *Synechocystis* host strain to be used as an *in vivo* screening tool where the activity of heterologously expressed *Gallionella* Rubisco, CbbM, affects the host’s growth rate. As a first step, we depleted endogenous Rubisco activity using CRISPRi (Fig. 1A). The genes encoding the Rubisco large and small subunits, *rbcL* and *rbcS*, are part of a polycistronic operon, *rbcLXS*, which is essential under photoautotrophic and heterotrophic growth conditions (11, 32). We used anhydrotetracycline (aTc)-inducible CRISPR interference (CRISPRi) to conditionally repress native Rubisco, RbcLS, production. The CRISPRi construct consisted of the gene encoding dCas9 and two single guide RNAs (sgRNAs), which were complementary to DNA stretches located within the first 70 bp of the *rbcL* coding sequence. A transcriptional repression of *rbcL* targets the whole *rbcLXS* operon. We used mass spectrometry to compare the depletion of RbcLS in strains harboring either only the gene for dCas9 and no sgRNA (*Syn*-sgRNA(−)) or the complete CRISPRi construct including the *rbcLXS*-specific sgRNAs (*Syn*-sgRNA*_rbc_*) after induction with aTc for approximately five generations (Fig. 1B, Supp. Fig. S1A and B, Supp. Tables S1 and S2). Only 1% of RbcL and 5% of RbcS were detected in *Syn*-sgRNA*_rbc_* compared to the system without target-specific sgRNAs in *Syn*-sgRNA(−). We observed a similar trend for a strain containing the complete CRISPRi system and complemented with wild-type *Gallionella* Rubiso, CbbM, expressed from a plasmid (*Syn*-sgRNA*_rbc_ cbbM*(WT)) (2% of RbcL, 3% of RbcS, Fig. 1B, Supp. Fig. S1A to D, Supp. Tables S1 to S4). The observed depletion matches our prior experiences of approximately 75% transcript repression using the CRISPRi system when targeting a variety of other genes, e.g. *glgC* (33), *gltA* (34) or *gyrA* (35).

**Figure 1:**
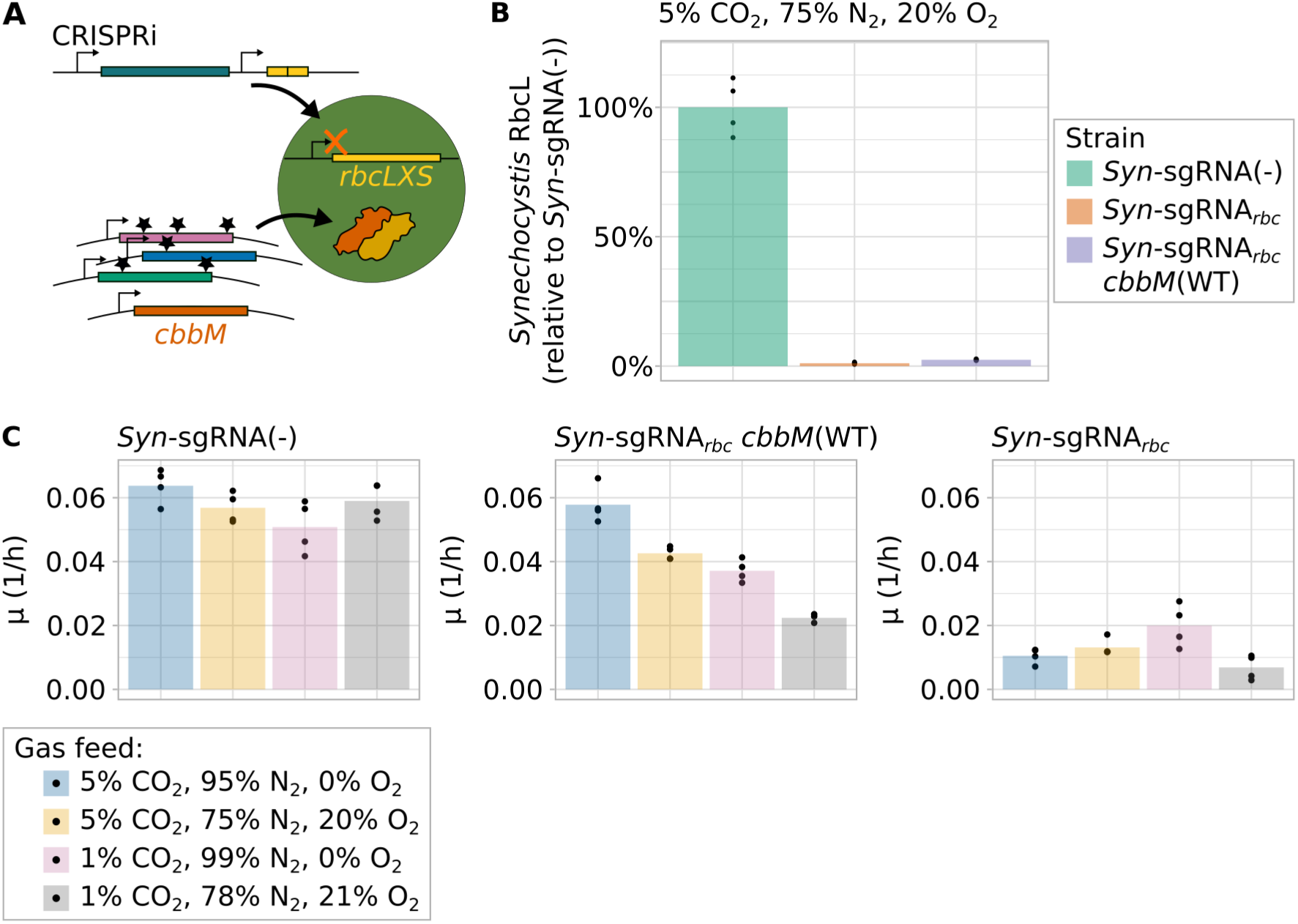
Characterization of the host strain used for Rubisco screening. (**A**) Schematic of method for screening of mutational variants. CRISPR inhibition (CRISPRi) was used for transcriptional repression of the *rbcLXS* transcript encoding the endogenous Rubisco of *Synechocystis*. The gene *cbbM*, which encodes *Gallionella* Rubisco, CbbM, was introduced into the host strain under the control of a constitutive promoter. (**B**) Relative amount of *Synechocystis* Rubisco large subunit RbcL measured by mass spectrometry comparing *Syn*-sgRNA(−), *Syn*-sgRNA*_rbc_*and *Syn*-sgRNA*_rbc_ cbbM*(WT) after approximately five generations of induction of the CRISPRi system using anhydrotetracycline (aTc). Signal intensity as measured by mass spectrometry for RbcL, the *Synechocystis* Rubisco large subunit, is shown relative to the value measured for *Syn*-sgRNA(−). Bar plots represent the mean of four replicates, which are shown as single dots. For non-normalized data and data for the small subunit RbcS, see Supp. Fig. S1. (**C**) Growth rates of *Syn*-sgRNA(−), *Syn*-sgRNA*_rbc_*and *Syn*-sgRNA*_rbc_ cbbM*(WT) in batch cultivation for different CO_2_/O_2_ ratios as gas feed after induction of the CRISPRi system using aTc (n=4, n=3 for *Syn*-sgRNA*_rbc_ cbbM*(WT) at 1% CO_2_, 78% N_2_, 21% O_2_). Cells were grown as described for panel (A) and then shifted to the indicated gas conditions. Growth was recorded after the shift to different gas conditions and back dilution to OD_730nm_ ∼ 0.1. For panels (B) and (C), all cultivations were performed at a light intensity of 300 µE. The corresponding growth curves are shown in Supplementary Figure 2.

### Growth rate of *Synechocystis* with CbbM is dependent on the CO_2_/O_2_ ratio in the gas feed

Contrasting to the *Synechocystis* Form I RbcLS, CbbM lacks the protective environment of carboxysomes. Carboxysomes are proteinaceous microcompartments which encapsulate Rubisco in close proximity to the enzyme carbonic anhydrase, and therefore help to enrich free CO_2_ around Rubisco and potentially lower the local O_2_ concentration (36). We confirmed the cytoplasmic localization of CbbM by microscopy (Supp. Fig. S2). Previous reports on the expression of the Type II Rubisco from *R. rubrum* also showed cytoplasmic localization (11).

For cytoplasmically localized CbbM, we assume that enzyme kinetic parameters play a large role in determining a strain’s growth rate in dependence on different gas feed compositions (compare Supp. Results & Discussion). To test this hypothesis, we recorded batch cultivation growth curves of *Syn*-sgRNA(−), *Syn*-sgRNA*_rbc_* and *Syn*-sgRNA*_rbc_ cbbM*(WT) at different gas feed compositions with varying CO_2_/O_2_ ratios and determined their growth rate in a time period briefly after changing to the gas feed of interest (Fig. 1C and Supp. Fig. S3). Under all tested conditions, *Syn*-sgRNA(−) grew the fastest of the three strains and *Syn*-sgRNA*_rbc_*the slowest (Fig. 1C, Supp. Fig. S3). Different gas feed compositions had no strong effect on the growth rates of *Syn*-sgRNA(−) or *Syn*-sgRNA*_rbc_*. Carboxysome encapsulation of RbcLS in *Syn*-sgRNA(−) likely protects the enzyme from changes in the exogenous gas feed. The slow growth of *Syn*-sgRNA*_rbc_*reflects the fact that RbcLS activity is essential for photoautotrophic growth of *Synechocystis* (11, 32). Residual growth might be explained by a low remaining amount of RbcLS in the cell (Fig. 1B).

At a high CO_2_/O_2_ ratio in the gas feed (5% CO_2_, 95% N_2_, 0% O_2_), expression of CbbM rescued the growth deficiency of *Syn*-sgRNA*_rbc_* and the strain *Syn*-sgRNA*_rbc_ cbbM*(WT) grew almost as fast as *Syn*-sgRNA(−) (Fig. 1C). Adding O_2_ (5% CO_2_, 75% N_2_, 20% O_2_) to the gas feed reduced the growth rate of *Syn*-sgRNA*_rbc_ cbbM*(WT), implying that the enzyme’s poor specificity value might play a role at this gas feed composition.

When reducing the CO_2_ concentration in the gas feed to 1% CO_2_, 99% N_2_, 0% O_2_, the growth deficiency of *Syn*-sgRNA*_rbc_ cbbM*(WT) was slightly exacerbated, likely reflecting CbbM operating below the limiting rate *V_C_* due to CO_2_ substrate limitation (compare Supp. Results & Discussion). Alternatively, the oxygen generation from photosynthesis might lead to a detrimental ratio of CO_2_ to O_2_, leading to RuBP oxygenation and therefore reduced growth. When adding oxygen to the 1% CO_2_ gas feed (1% CO_2_, 78% N_2_, 21% O_2_), the growth rate of *Syn*-sgRNA*_rbc_ cbbM*(WT) was reduced markedly, almost to that of the CRISPRi strain with no CbbM and only residual RbcLS activity. Thus, CbbM cannot support growth in a condition with 1% CO_2_, 78% N_2_, and 21% O_2_. The growth rates of *Syn*-sgRNA*_rbc_ cbbM*(WT) at different gas feed compositions are in line with those reported for a strain in which *Synechocystis* RbcLS was exchanged by *R. rubrum* Form II Rubisco (11).

By changing the gas mixture, our method can be used to select for improvements in different catalytic properties. At a high CO_2_/O_2_ ratio in the gas feed (5% CO_2_, 95% N_2_, 0% O_2_), we assume that the created host strain can be used as a chassis to screen for variants with a changed *V_C_*. When lowering the CO_2_/O_2_ ratio, e.g. by using a gas feed of 5% CO_2_, 75% N_2_ and 20% O_2_, the specificity of a Rubisco variant for CO_2_ over O_2_, S(℅), should play a more decisive role in determining a strain’s growth rate, as the oxygenation reaction directs RuBP substrate away from the growth-benefitting carboxylation reaction. When lowering the substrate concentration to 1% CO_2_, the enzyme’s affinity for CO_2_, expressed by *K_C_*, should become more important for the growth rate.

Unlike carboxysome-located RbcLS, CbbM is dependent on cytosolic free CO_2_ and might be substrate-limited even in a gas feed containing 5% CO_2_. We explored a strategy to increase the intracellular concentration of CO_2_ by deleting the NDH-1 subunits NdhD3 and NdhD4, encoded by genes *ndhD3* (*sll1733*) and *ndhD4* (*sll0027*), which abolishes the rapid transformation of cytosolic free CO_2_ into carbonate by the NDH-1 complex (36, 37). The deletion did not increase the growth rate of *Syn*-sgRNA*_rbc_ cbbM*(WT) Δ*ndhD3* Δ*ndhD4* compared to *Syn*-sgRNA*_rbc_ cbbM*(WT) at gas feeds of 1% or 5% CO_2_ without added O_2_ (Supp. Fig. S4). Neither did increasing the CO_2_ concentration in the gas feed to 10% (10% CO_2_, 90% N_2_, 0% O_2_, data not shown). It has to be noted that a gas feed of 10% CO_2_ might be inhibitory to *Synechocystis* growth, as judged by the effect of high CO_2_ feeds on *Cyanothece* 51142 and *Synechococcus elongatus* PCC 11801 (38, 39). Alternative strategies to increase the strain’s growth rate are introducing the putative CbbM-specific Rubisco activase (40) or by engineering a co-localization of CbbM and carbonic anhydrase, for instance in a carboxysome(-like) structure.

### Proteomics shows stress signatures when replacing RbcLS with CbbM

To investigate the effect of RbcLS depletion and CbbM expression on the cells, we performed untargeted label-free DIA proteomics on the different strains when grown at 5% CO_2_, 75% N_2_, 20% O_2_ approximately five generations after addition of aTc (Fig. 2A and B, Supp. Fig. S5A, Supp. Tables S1 and S2). We unambiguously identified 1,873 proteins. When comparing the strain *Syn*-sgRNA*_rbc_ cbbM*(WT) at different gas feeds (Fig. 2C and D, Supp. Fig. S5B, Supp. Tables S3 and S4), we unambiguously detected 1,857 proteins. Both numbers correspond to a coverage of approximately 52% of annotated coding sequences.

**Figure 2:**
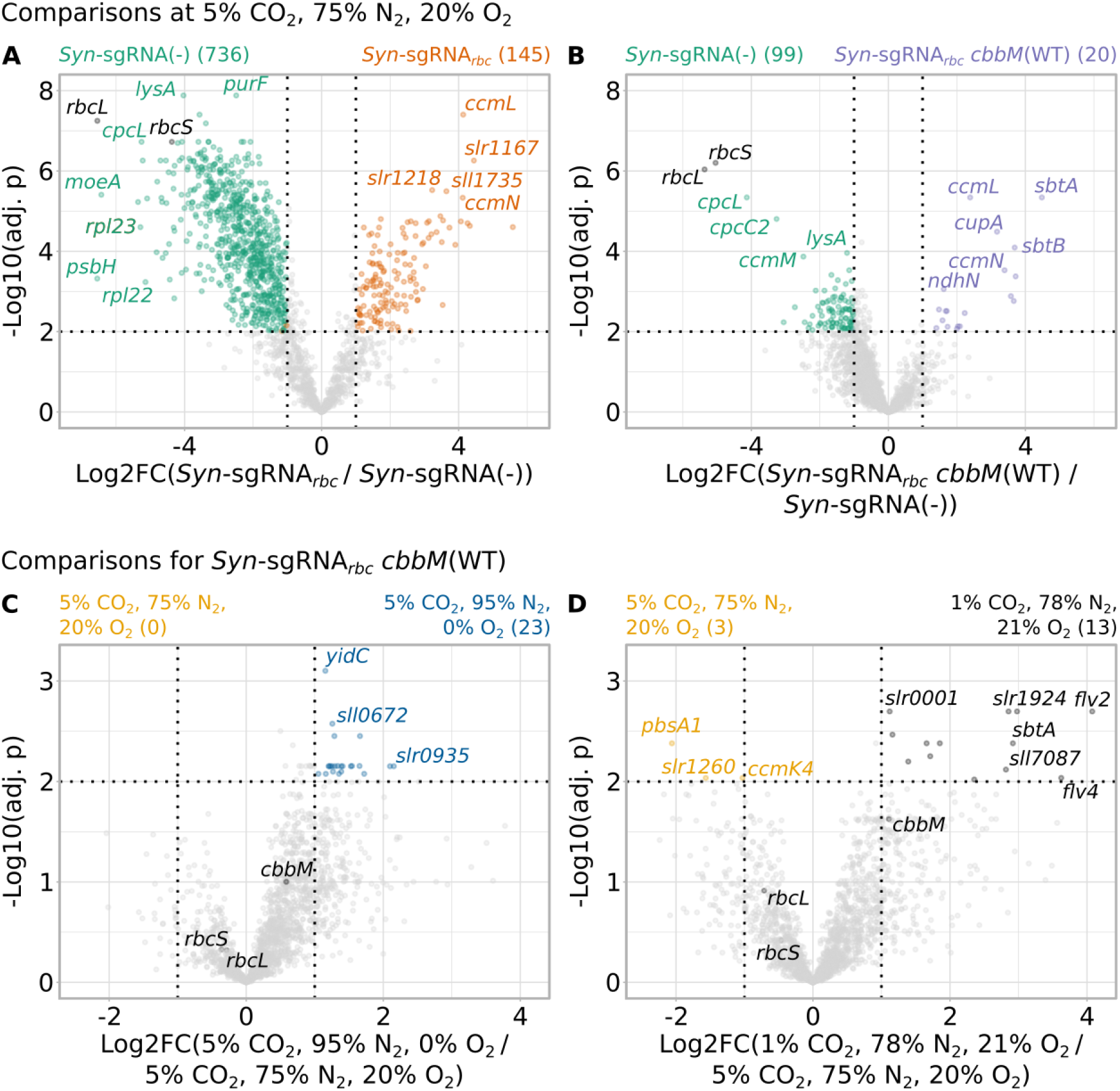
Untargeted proteomics comparing the proteome of different strains at a gas feed of 5% CO_2_, 75% N_2_, 20% O_2_ (panels **A** and **B**) and of *Syn*-sgRNA*_rbc_ cbbM*(WT) at different gas feed compositions (panels **C** and **D**). Samples were taken as described for Fig. 1B and untargeted proteomics was performed. Numbers in parentheses indicate the number of proteins upregulated in the respective strain or condition with *|Log2FC| > 1* and *adj. p < 0.01* and dotted lines indicate these respective cut-off values.

Knocking down RbcLS in *Syn-*sgRNA*_rbc_* had a more severe effect on the proteome compared to *Syn*-sgRNA(−) than replacing it by CbbM in *Syn*-sgRNA*_rbc_ cbbM*(WT) (Fig. 2A and B), which matches the relative growth rate differences of the strains at 5% CO_2_, 75% N_2_, 20% O_2_ (Fig. 1C). Fewer gene ontology (GO) and KEGG pathways were affected when comparing *Syn*-sgRNA(−) to *Syn*-sgRNA*_rbc_ cbbM*(WT) than to *Syn-*sgRNA*_rbc_* (Supp. Fig. S6 and S7, Supp. Tables S5–S8), which underlines that expression of CbbM can rescue the effect of the depletion of RbcLS to a certain extent. Still, there seems to be a certain degree of metabolic stress on the cell based on the fact that several central biological processes were down-regulated in *Syn*-sgRNA*_rbc_ cbbM*(WT) compared to *Syn*-sgRNA(−).

The first step in carboxysome assembly is the interaction between RbcLS, carbonic anhydrase and CcmM (36). The lack of its interaction partners *Synechocystis* RbcL and RbcS seemed to lead to a destabilization of CcmM in both *Syn*-sgRNA*_rbc_* and *Syn*-sgRNA*_rbc_ cbbM*(WT) (Table 1, underlining the loss of carboxysome structures in both strains.

**Table 1:** Subset of Supplementary Tables S1 and S2 with data for the Rubisco operon, carboxysome, carbonic anhydrase, CmpR and NdhR regulon according to (36). Comparisons are given for different strains at a gas feed of 5% CO_2_, 75% N_2_, 20% O_2_ (compare Fig. 2, Supp. Fig. S5). “ND” signifies that the respective number could not be determined, e.g because the protein of interest was not detected in one of the samples. Log2 fold changes (Log2FC) above 1.0 or below −1.0 with associated significant adjusted p values (*adj. p < 0.01*) were marked in green and red, respectively.

|  |  |  | Syn-sgRNA<sub>rbc</sub>/<br>Syn-sgRNA(-) |  | Syn-sgRNA<sub>rbc</sub> cbbM(WT)/<br>Syn-sgRNA(-) |  |
| --- | --- | --- | --- | --- | --- | --- |
|  | Locus tag | Gene name | Log2FC | adj. p | Log2FC | adj. p |
| Rubisco | *slr0009* | *rbcL* | -6.55 | 5.7E-8 | -5.37 | 9.1E-7 |
| Rubisco | *slr0012* | *rbcS* | -4.37 | 1.9E-7 | -5.04 | 6.3E-7 |
| Carboxysome | *sll1028* | *ccmK2* | 3.47 | 1.3E-5 | 0.78 | 1.2E-1 |
| Carboxysome | *sll1029* | *ccmK1* | 0.70 | 4.1E-1 | 0.49 | 6.7E-1 |
| Carboxysome | *sll1030* | *ccmL* | 4.13 | 4.0E-8 | 2.39 | 4.6E-6 |
| Carboxysome | *sll1031* | *ccmM* | -2.64 | 5.1E-6 | -2.48 | 1.4E-4 |
| Carboxysome | *sll1032* | *ccmN* | 4.12 | 4.6E-6 | 3.39 | 3.0E-4 |
| Carboxysome | *slr0169* | *ccmP* | -3.40 | 4.4E-6 | -0.35 | 3.7E-1 |
| Carboxysome | *slr0436* | *ccmO* | 0.59 | 1.0E-1 | 1.05 | 3.6E-2 |
| Carboxysome | *slr1838* | *ccmK3* | -0.04 | 9.0E-1 | -0.60 | 1.1E-1 |
| Carboxysome | *slr1839* | *ccmK4* | -1.90 | 2.3E-5 | -0.56 | 6.8E-2 |
| Carbonic anhydrase | *slr1347* | *ccaA* | -1.18 | 3.0E-2 | -1.59 | 2.7E-2 |
| CmpR regulon | *slr0042* | *slr0042* | -1.11 | 3.0E-2 | 0.13 | 8.5E-1 |
| CmpR regulon | *slr0043* | *cmpC* | ND | ND | -0.55 | 2.0E-1 |
| NdhR regulon | *sll1594* | *ndhR* | -0.01 | 9.8E-1 | 0.85 | 4.1E-2 |
| NdhR regulon | *slr2011* | *slr2011* | 1.02 | 2.0E-2 | 0.21 | 6.8E-1 |
| NdhR regulon | *sll1733* | *ndhD3* | ND | ND | 3.66 | 1.7E-3 |
| NdhR regulon | *sll1734* | *cupA* | 1.70 | 1.0E-4 | 3.17 | 3.2E-5 |
| NdhR regulon | *sll1735* | *sll1735* | 3.64 | 3.2E-6 | 1.71 | 3.1E-3 |
| NdhR regulon | *slr1512* | *sbtA* | 0.80 | 1.2E-2 | 4.48 | 4.6E-6 |
| NdhR regulon | *slr1513* | *sbtB* | 2.22 | 1.1E-4 | 3.69 | 8.0E-5 |
| NdhR regulon | *sll0529* | *sll0529* | -2.76 | 7.4E-7 | -0.13 | 6.0E-1 |

### Reduced carboxylation and/or increased RuBP oxygenation in sgRNA*_rbc_ cbbM*(WT)

*Synechocystis* cells sense low carbon availability by integrating the levels of different metabolites such as 2PG, RuBP or 2-oxoglutarate (2OG) (36). For instance, the repressing transcription factor NdhR is induced by 2OG and repressed by 2PG (36). The metabolites RuBP and 2PG induce the activating transcription factor CmpR (36). In *Syn*-sgRNA*_rbc_*, and to an even higher extent in *Syn*-sgRNA*_rbc_ cbbM*(WT), several proteins belonging to the NdhR regulon were significantly more abundant than in *Syn*-sgRNA(−) (Table 1). Interestingly, levels of proteins belonging to the CmpR regulon did not change significantly in the CRISPRi strains in comparison to *Syn*-sgRNA(−).

As described above, CbbM lacks the protective environment of the carboxysome (Supp. Fig. S2) and is thus not colocalized with carbonic anhydrase. This should lead to a decreased CO_2_/O_2_ ratio in the immediate surroundings of CbbM compared to carboxysome-encapsulated RbcLS (compare (41)). As a consequence, we might expect reduced carboxylation and thus a lowered 2OG level and/or an elevated 2PG level due to increased RuBP oxygenation in *Syn*-sgRNA*_rbc_ cbbM*(WT) compared to *Syn*-sgRNA(−).

In *Syn*-sgRNA*_rbc_*, remaining RbcLS should be carboxysome-localized. Proteomic changes should therefore not be triggered by RuBP oxygenation, but reduced RuBP carboxylation due to limiting Rubisco activity. Hence, both 2PG and 2OG levels should drop in comparison to *Syn*-sgRNA(−). Induction of the NdhR regulon in *Syn*-sgRNA*_rbc_*is thus probably only the result of lower carbon fixation.

Proteins belonging to the NdhR regulon were more abundant in *Syn*-sgRNA*_rbc_ cbbM*(WT) than in *Syn*-sgRNA*_rbc_*, hinting towards increased RuBP oxygenation in *Syn*-sgRNA*_rbc_ cbbM*(WT) (Table 1). Despite showing a similar high-carbon requiring phenotype, proteomic changes of *Syn*-sgRNA*_rbc_ cbbM*(WT) at 5% CO_2_, 75% N_2_, 20% O_2_ did not resemble the transcriptomic changes found for a carboxysome-less *Synechocystis* strain lacking the main constituent of the carboxysome, *ccmM*, when grown at 5% CO_2_, ambient O_2_ (41).

### Photosynthetic oxygen production likely sufficient to cause increased photorespiration in *Syn*-sgRNA*_rbc_ cbbM*(WT)

In a second step, we compared the proteome of *Syn*-sgRNA*_rbc_ cbbM*(WT) at different gas conditions (Fig. 2C and D, Supp. Fig. S5B). On a proteomic level, we did not observe clear evidence for the underlying reason for the strongly reduced growth rate of *Syn*-sgRNA*_rbc_ cbbM*(WT) at 1% CO_2_ compared to growth at 5% CO_2_. GSEAs comparing the proteome at gas feeds with 5% CO_2_ or 1% CO_2_ and added oxygen did not identify any KEGG terms or GO terms associated with biological processes as enriched or depleted (Supp. Table S13). Proteomic changes of *Syn*-sgRNA*_rbc_ cbbM*(WT) at 1% CO_2_, 78% N_2_, 21% O_2_ (Supp. Table S4) were reminiscent of transcriptomic changes of long-term low-CO_2_ acclimated wild-type *Synechocystis*, e.g. an upregulation of SbtA, which belongs to the NdhR regulon, and of the *flv2*–*flv4* operon (41, 42).

When comparing *Syn*-sgRNA*_rbc_ cbbM*(WT) grown in a gas feed of 5% CO_2_ with or without O_2_, we did not observe a significant reaction of proteins belonging to the NdhR or CmpR regulons (Supp. Table S9), even though RuBP oxygenation and thus 2PG production should be higher in the oxygen-containing than in the oxygen-free gas feed. Hence, based on the relative increase of proteins belonging to the NhdR regulon in *Syn*-sgRNA*_rbc_ cbbM*(WT) at both gas feeds compared to *Syn*-sgRNA(−) (Table 1), it can be assumed that either the endogenously produced oxygen was sufficient to cause increased photorespiration in the oxygen-free gas feed or that a relative decrease of the local CO_2_ concentration in immediate vicinity to the enzyme compared to carboxysome-localized RbcLS lead to decreased carboxylation and lowered 2OG levels.

The set of 23 proteins with significantly higher levels in the culture grown without oxygen in the gas feed (*|Log2FC| > 1.0*, *adj. p < 0.01*, Supp. Table S10) did not give clear evidence on the differences in the metabolic states of the cells in the two different gas feed conditions. Compared to the aerobic gas feed, proteins belonging to the GO term “photosynthesis” and KEGG pathways “Photosynthesis” as well as “ABC transporters” were enriched in the cells grown in the anaerobic gas feed (Supp. Fig. S7, Supp. Tables S11 and S12). The “ABC transporters” reacting most strongly encompassed phosphate, amino acid and urea transporters, which might reflect the higher growth rate under this condition (Fig. 1C).

### Phylogenetically-backed strategy to predict beneficial higher-order variants

In a next step, we wanted to test if the established host strain can be used to find amino acid replacements that enhance Rubisco activity by performing growth competition experiments of pooled *Gallionella* CbbM variant libraries. A well designed input library, a “zero-shot prediction”, can help to explore protein fitness landscapes more efficiently than random input if there is no experimental data on protein mutants available (23). Additionally, libraries of higher-order mutational variants are better suited for exploring the protein fitness landscape of a protein than single-site mutational variants since they provide valuable information about epistasis (43–45). This might be especially important for Rubisco. A recent DMS of the complete Form II *R. rubrum* Rubisco did not identify any single amino acid exchange with a strong beneficial impact on the catalytic constant of the enzyme (17).

However, the chance of losing enzyme functionality increases with the extent of amino acid exchanges (18–22). Furthermore, prior studies attempting to rationally engineer Rubisco based on, for instance mechanistic assumptions, showed that this is a challenging endeavor (1, 2). Hence, we used phylogenetic information to predict the fitness of enzyme variants using the bioinformatic tool EVmutation (24, 25). EVmutation uses multiple sequence alignments of homologues of the protein of interest to calculate the conservation of specific amino acids at each position, which results in an independent prediction score (24, 25). It enriches this information by calculating the pairwise co-occurrence of amino acids at different positions, i.e. their “evolutionary coupling” or also “direct coupling”, leading to an “epistatic” prediction score.

For our test library, we decided to combine five beneficial amino acid exchanges in a combinatorial manner and hence a library of in total 2^5^ = 32 different variants, because this number of variants is experimentally easily tractable. To select five mutations from the many possibilities, we used *in silico* evolution, which is a variant of the Metropolis-Hastings algorithm (27) to heuristically explore the protein fitness landscape given by the EVmutation model (Methods). Based on occurrence and co-occurrence frequencies during the *in silico* evolution, as well as predicted fitness values of the combinatorial variants, we decided on exchanges M140E, S162P, A230E, V323A and G422R (Fig. 3B and C, Table 2, Supp. Table 15). These were also among the ten replacements with the highest predicted fitness when introduced on their own (Supp. Table S14) and, according to EVmutation, not directly coupled among each other (Table 2, Supp. Fig. S9). Indeed, the residues are located in different parts of the protein structure (Fig. 3C). None of the amino acids was in close proximity to the enzyme’s catalytic center (roughly depicted by K200, a highly conserved lysine which is carbamylated during and essential for catalysis, shown in black in Fig. 3C). Furthermore, all suggested residues faced away from the enzyme’s dimerization interface. Exchanges should therefore neither affect dimerization nor RuBP binding directly. All suggested exchanges except V323A were surface-exposed.

**Figure 3:**
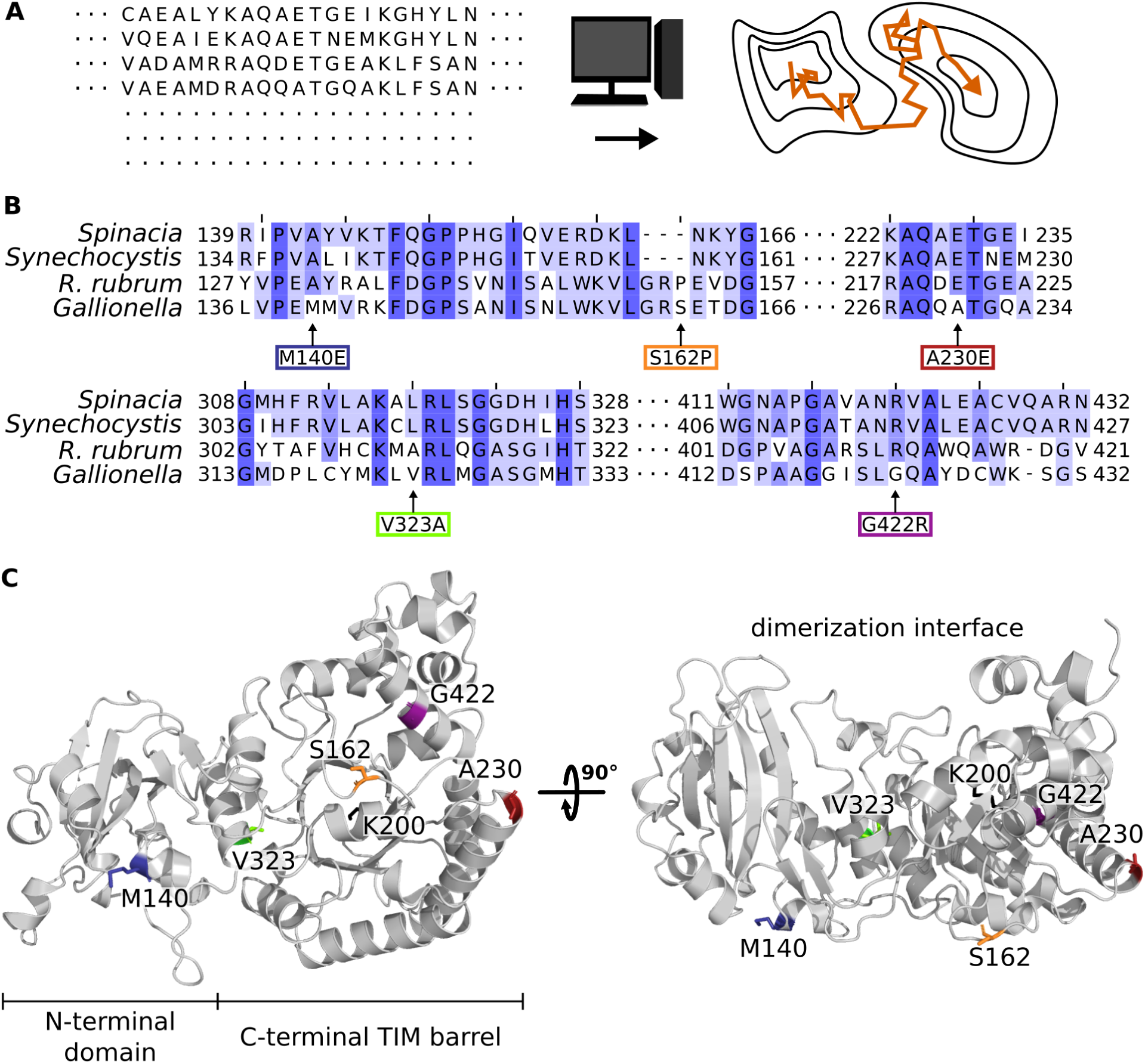
Phylogenetically-backed strategy to predict beneficial amino acid exchanges. (**A**) A multiple sequence alignment for *Gallionella* Rubisco was created and used as input for the EVcouplings web server. The resulting model was used to perform *in silico* evolution and identify new local fitness maxima in order to suggest beneficial *cbbM* variants. *In silico* evolution describes a heuristic algorithm to identify local fitness maxima by traversing “fitness valleys” between different “fitness maximum peaks”. (**B**) Alignment of fragments of the amino acid sequences of several Rubisco homologues, specifically of *Spinacia oleracea* (*Spinacia*, P00875), *Synechocystis* sp. PCC 6803 (*Synechocystis*, P54205), *Rhodospirillum rubrum* (*R. rubrum*, P04718), *Gallionella* sp. (*Gallionella*, A0A1G0AW29). Arrows point at the amino acid exchanges highlighted in (C). Shades of blue indicate different degrees of identity between the compared protein sequences. Numbers give the amino acid position of the first and last amino acid shown in the alignment. (**C**) Structure of *Gallionella* Rubisco as cartoon model, on the left shown from the side which is facing away from the dimerization interface. On the right, the structure is shown turned by 90°. Amino acid positions suggested to be mutated and carbamylated lysine K200 are highlighted in color and shown as stick models.

**Table 2:**
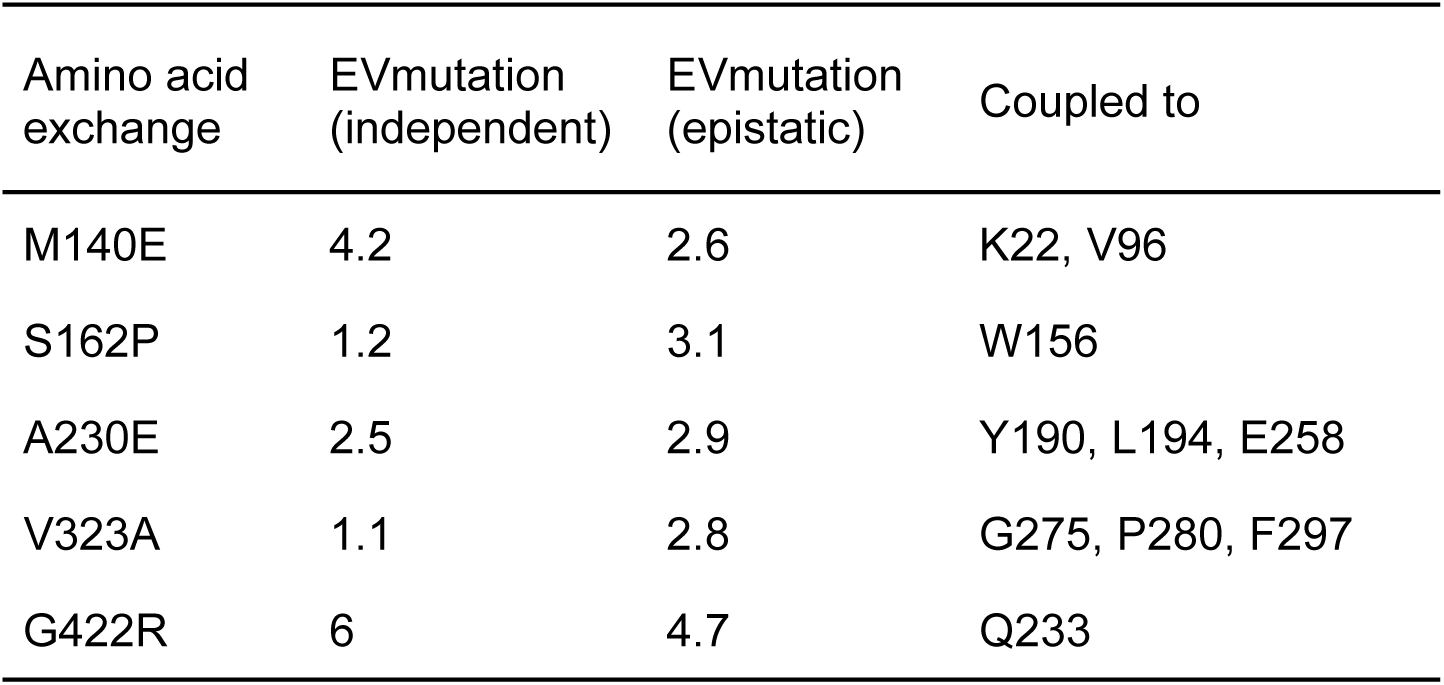
Amino acid exchanges suggested by EVmutation coupled to Metropolis-Hastings algorithm. EVmutation independent and epistatic fitness predictions are given. Epistatic fitness predictions take evolutionary couplings into account, whereas independent scores are only based on occurrence frequencies. Column “coupled to” gives residues that are predicted to be evolutionary coupled to the respective position. For fitness predictions of all possible amino acid exchanges covered by the used MSA compare Supp. Table S14.

| Amino acid exchange | EVmutation (independent) | EVmutation (epistatic) | Coupled to |
| --- | --- | --- | --- |
| M140E | 4.2 | 2.6 | K22, V96 |
| S162P | 1.2 | 3.1 | W156 |
| A230E | 2.5 | 2.9 | Y190, L194, E258 |
| V323A | 1.1 | 2.8 | G275, P280, F297 |
| G422R | 6 | 4.7 | Q233 |

It is noteworthy that all suggested exchanges but M140E changed amino acids to the one present in the *R. rubrum* Rubisco background (Fig. 3B, *R. rubrum* amino acid/*Gallionella* amino acid: P153/S162, E221/A230, A312/V323, R411/G422). Amino acid replacements may show sign epistasis, i.e. might show opposite effects depending on the remaining sequence background (19, 43, 44). Nevertheless, we checked the approximate effects of exchanges at the suggested positions according to a recently published DMS of *R. rubrum* Rubisco (17). Exchanges at homologous sites of the suggested positions did not show a strong effect on catalytic parameters in *R. rubrum* Rubisco (*R. rubrum* amino acid/*Gallionella* amino acid: P153/S162, E221/A230, A312/V323, R411/G422) with the exception of A312/V323 (*R. rubrum*/*Gallionella* position).

In a next step, we cloned the respective *cbbM* gene variants and introduced them into *Syn*-sgRNA*_rbc_*. We were not able to obtain any *E. coli* colonies for constructs into which we tried to introduce amino acid exchange M140E. Hence, we continued with a 16-member library based on the remaining four amino acid exchanges. The EVmutation fitness values for these 16 variants are given in Supp. Table S15.

### *In vivo* screening of CbbM variants at different growth conditions

We performed a pooled growth competition assay of this 16-member library (Fig. 4A). Each of the strains harbored a 20 nucleotide long strain-specific barcode at the 3’ end of the *cbbM* gene (not translated), which enabled us to track each strain’s relative abundance in the pool by next generation sequencing (Fig. 4A). We grew the pool of the 16 strains in a photobioreactor in turbidostat mode, i.e. pooled cultures were automatically back-diluted to an OD_720nm_ of approximately 0.2. These regular dilutions resulted in strains with lower growth rates being diluted out of the liquid culture over time and variants with faster growth rates accumulating. By sequencing the barcodes present in the liquid culture over time, the relative abundance of the different strains were tracked (Fig. 4B, Supp. Fig. S10, S11 and S12, Supp. Table S16). These relative abundances were used to calculate fitness values associated with each strain and growth condition (Fig. 4C, Supp. Table S16).

**Figure 4:**
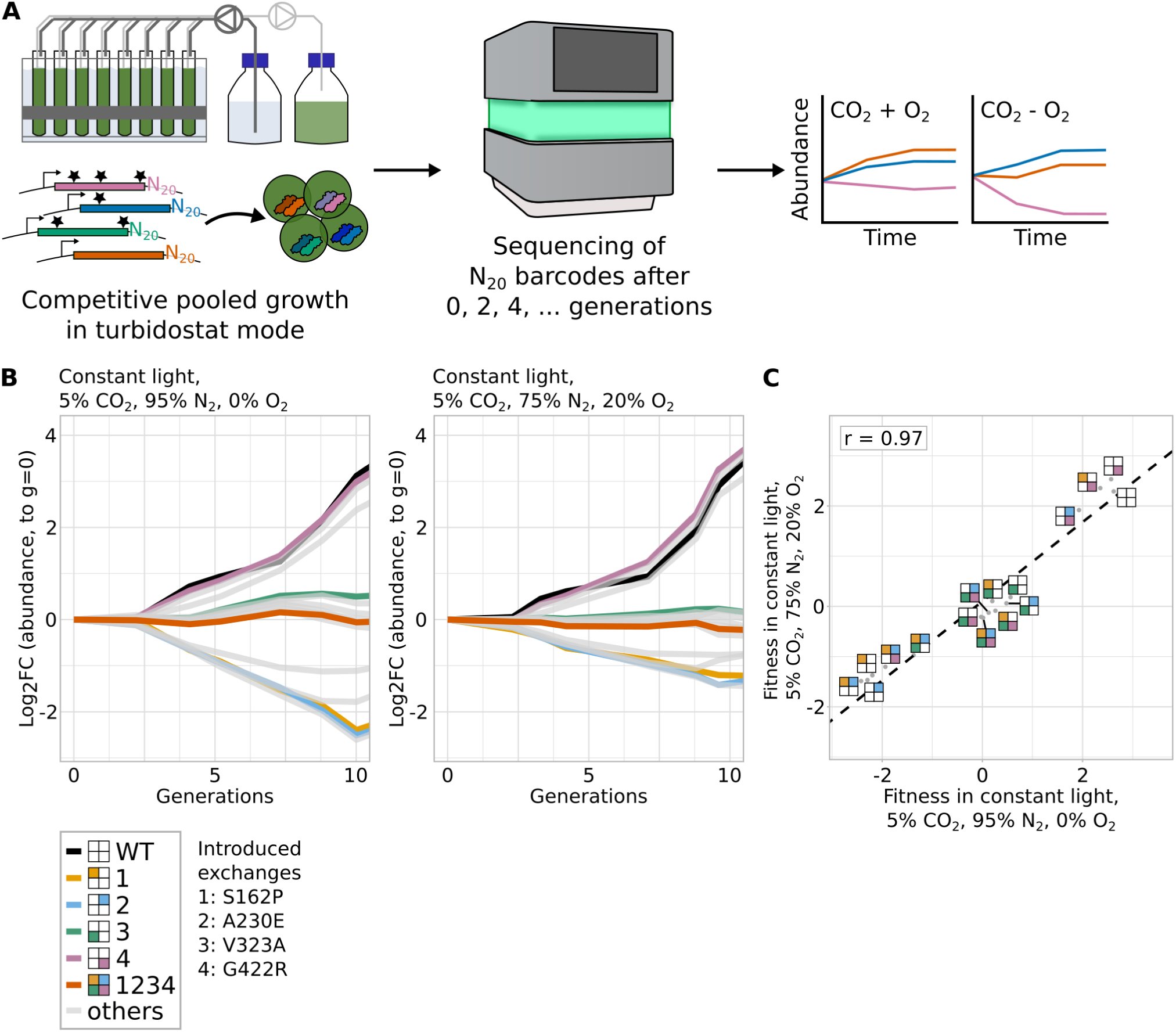
*In vivo* screening of library of Rubisco variants. (**A**) Scheme illustrating high-throughput screening by competitive growth of the pooled variant library (left) followed by next generation sequencing (middle) and fitness calculations (right). (**B**) Changes of abundance of different strains during competitive growth of the 16-member library at different growth conditions. For reasons of clarity, data of only the wild-type variant, variants with a single amino acid exchange and of the quadruple mutant variant are depicted in color. Fitness values of other variants are plotted in light gray. For a plot including 95% confidence intervals of the highlighted variants, see Supp. Fig. S11A. Fitness data of all variants is shown in Supp. Fig. S11B. Light intensity for all cultivations was 300 µE. (**C**) Scatter plots comparing the fitness data from different growth conditions. The Pearson correlation coefficient r is given for the comparison of fitness values at the two different continuous light conditions (*r = 0.97, p = 1.833e-10*). Panels (B) and (C): For reasons of clarity, we depict every variant as consisting of four boxes which represent the four different introduced amino acid exchanges. If a box is colored, the exchange was introduced and if left white, the exchange is absent.

We performed growth competition assays at different gas feeds and light conditions. Based on our pre-experiments (Fig. 1C), we included growth at continuous light of 300 µE and a gas feed of 5% CO_2_ with or without added oxygen (5% CO_2_, 95% N_2_, 0% O_2_ and 5% CO_2_, 75% N_2_, 20% O_2_). For both conditions, we ran four replicate cultivations, which were inoculated from the same mutant pool.

The different CbbM variants were detected in similar amounts prior to the start of the growth competition experiments (gini for all samples of the original pool < 0.15). The replicates of both conditions showed a high correlation per time point, as measured by Pearson’s r (r > 0.8 among replicates for 80% of samples).

Fitness values of different mutant variants were highly correlated between both continuous light growth conditions (Fig. 4C), implying that none of the variants changed CbbM specificity to a relevant degree. In continuous light, no mutant variant gave a significantly higher fitness value than the wild-type enzyme with three variants being on par with it.

### *In vivo* pooled screening identified epistasis among amino acid exchanges

In the growth competition experiments described above, fitness values of different mutants separated into three groups of variants according to their relative fitness (Fig. 4B and C). Their fitness distribution revealed a small epistatic network among the tested amino acid exchanges (Fig. 4C, Fig. 5A). Exchanges S162P and A230E were detrimental when introduced on their own or when outnumbering V323A and G422R. The latter two rescued their negative impact. Furthermore, V323A was epistatic to G422R. In the following, to make naming of higher-order mutants easier, we will refer to the different replacements by numbers based on their relative position in the protein’s coding sequence: 1: S162P, 2: A230E, 3: V323A, 4: G422R. For instance, we will refer to the variant with S162P, V323A and G422R as CbbM(134), not CbbM(S162P, V323A, G422R). Note that amino acid numbering refers to the wild-type CbbM sequence excluding the 14 amino acid long Strep tag.

**Figure 5:**
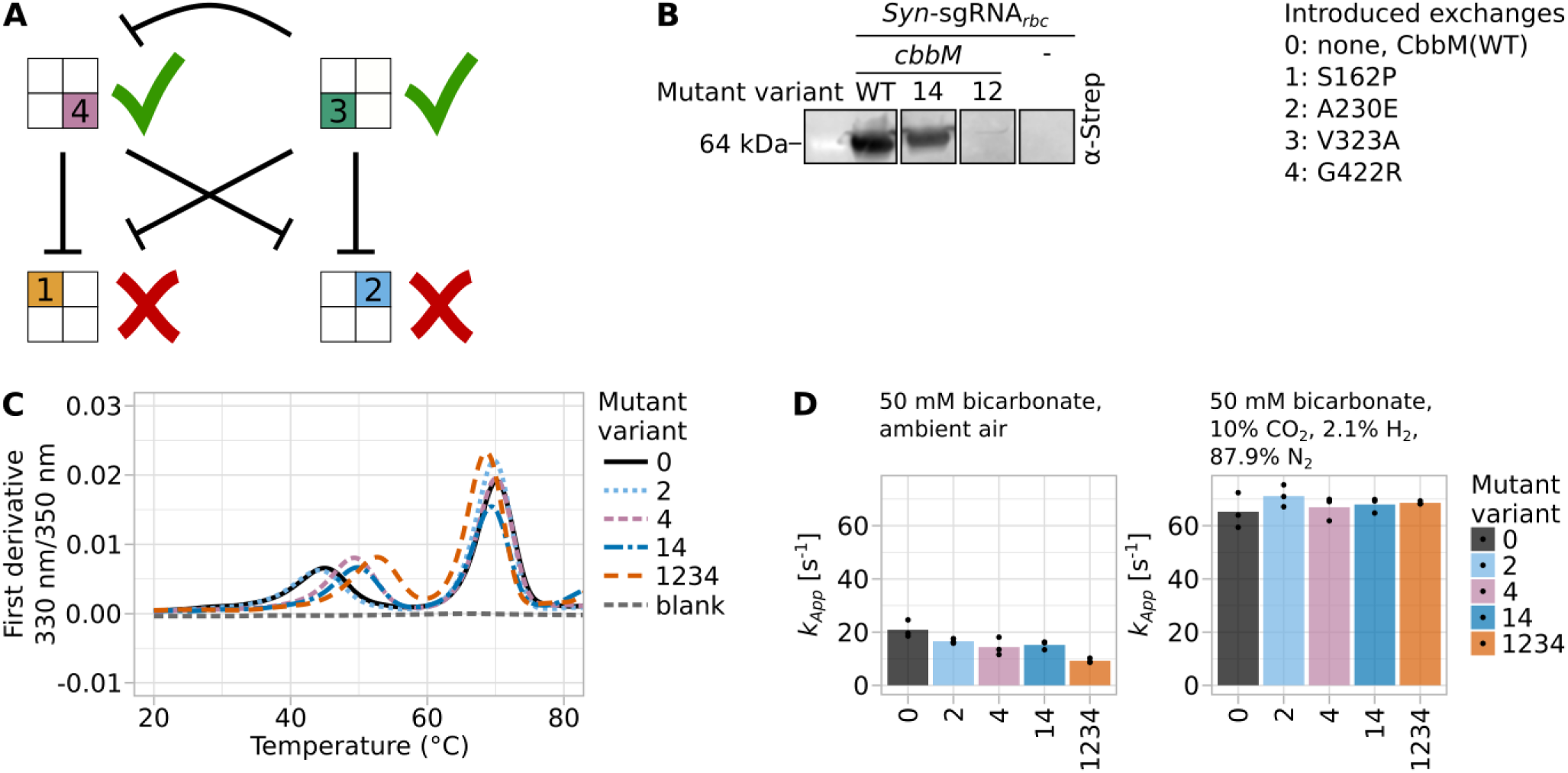
Epistasis among introduced amino acid exchanges. In all panels, introduced exchanges are coded as 1: S162P, 2: A230E, 3: S162P, 4: G422R. (**A**) Scheme illustrating epistasis of different mutations. For reasons of clarity, we depict every variant as consisting of four boxes which represent the four different introduced amino acid exchanges. If a box is colored, the amino acid exchange was introduced. If left white, the amino acid exchange is absent. (**B**) Western blot analysis of the soluble fraction of cell lysates of strains expressing selected mutant variants to determine the mutant protein’s abundance. For staining, horseradish peroxidase (HRP)-conjugated antibody directed against Strep tag (ɑStrep) was used. HRP chemiluminescence signal is overlaid with an image of the membrane. See Supp. Fig. S14 for the complete blot and, as a loading control, a Coomassie-stained SDS-PAGE gel loaded in the same manner as the gel used for the blot. The protein of interest, N-Strep-tagged CbbM, has a size of 53 kDa. (**C**) Thermal melting curves of several CbbM variants as determined by nanoDSF (compare Supp. Table S17). (**D**) Apparent rate constants as measured by spectrophotometric *in vitro* assays. The assays were conducted either in ambient air (approx. 0.04% CO_2_, 78% N_2_, 21% O_2_) or in a gas atmosphere without oxygen (10% CO_2_, 2.1% H_2_, 87.9% N_2_, 0% O_2_). In both cases, 50 mM bicarbonate was added to the reaction mixture.

Amino acid exchanges S162P and A230E were detrimental under all tested conditions (Fig. 4C). The group of variants that contained only these exchanges or, in addition to both, either V323A or G422R, grew the slowest. Accordingly, the growth rate of variant *Syn*-sgRNA*_rbc_ cbbM*(12), when grown on its own, was almost as low as the one measured for *Syn*-sgRNA*_rbc_*, the strain lacking *cbbM* complementation (Fig. 1C, Supp. Fig. S3, Supp. Fig. S13). Based on western blot analysis of these variants (Fig. 5B), this growth deficiency was probably based on a strongly reduced CbbM expression of variant CbbM(12) compared to the wild-type variant CbbM(WT) or variant CbbM(14). Hence, replacements S162P and/or A230E had a negative effect on CbbM expressibility, solubility or stability. All variants which included V323A, and which were not part of the first described group, formed a group which grew approximately at the pool’s average growth rate (Fig. 4B and C). The last group contained wild-type CbbM (strain *Syn*-sgRNA*_rbc_ cbbM*(WT)) as well as strains containing G422R and either S162P or A230E, but not both. These four strains outperformed all other strains (Fig. 4B and C). We confirmed the comparable growth rates of *Syn*-sgRNA*_rbc_ cbbM*(14) to *Syn*-sgRNA*_rbc_ cbbM*(WT) in axenic cultivations (Supp. Fig. S13). According to western blot analysis, soluble CbbM levels in both strains were very similar (Fig. 5B, Supp. Fig. S14). Hence, G422R seemed to rescue the detrimental effects of S162P and A230E, but had no positive effect when introduced on its own. Interestingly, introducing V323A in these variants made strains behave like the strains belonging to the second group, showing that V323A was epistatic to G422R (Fig. 5A).

### Batch growth confirms pooled growth

We confirmed the growth differences observed in the pooled growth experiment for strains *Syn*-sgRNA*_rbc_ cbbM*(WT), *Syn*-sgRNA*_rbc_ cbbM*(14) and *Syn*-sgRNA*_rbc_ cbbM*(12) when grown separately for continuous light and a gas feed with or without O_2_ (Supp. Fig. S13). As observed in the pooled growth experiment, variants with wild-type CbbM(WT) and variant CbbM(14) had a comparable growth rate, whereas the variant expressing CbbM(12) grew only somewhat faster than the negative control (Fig. 1C, Supp. Fig. S3, Supp. Fig. S13). Hence, pooling different mutant strains did not seem to affect relative growth advantages and the pooled growth approach enabled us to separate strains according to their ability to promote *Synechocystis* growth.

### Long-distance interactions between tested amino acid positions

Fitness as measured in our system did neither correlate with the predicted values given by EVmutation (Supp. Fig. S15A) nor with the total number of exchanges introduced compared to the wild-type variant (Supp. Fig. S15B). Contrasting to EVmutation predictions (Table 2), and despite their spatial separation in the enzyme, the investigated amino acid exchanges showed epistasis among each other. Similar long-range amino acid interactions were also found for other proteins (43, 45, 46) and also specifically Rubisco homologues (6, 47–49). EVmutation captures evolutionary pressures on the enzyme, which does not necessarily correlate with the pressures exerted in the concrete experimental system under investigation. Alternatively, correlation between EVmutation predictions and fitness values might only be observable when investigating a larger set of variants. All suggested exchanges but M140E suggested replacements to the *R. rubrum* Rubisco background (Fig. 3B). A possible explanation might be biases present in the sequence database used for the construction of the multiple sequence alignment (50), which might reduce the predictive power of EVmutation and similar algorithms. This highlights the need for further metagenomic screenings and the characterization of phylogenetically more diverse variants to capture the full diversity and potential of Rubisco (compare (10)).

### *In vitro* characterization hints towards CbbM expression and stability changes underlying *in vivo* growth differences

We characterized a small subset of the pooled library *in vitro*. The melting temperature of several variants was measured using nanoDFS and showed two distinct melting points for all investigated variants (Fig. 5C, Supp. Fig. S16, Supp. Table S17). The melting points of CbbM(WT) were approximately 45°C and 70°C. These were only minimally affected by the introduction of A230E in variant CbbM(2). The lower melting point increased to approximately 49°C for variants CbbM(4) and CbbM(14), hinting towards a stabilizing effect of exchange G422R. This was raised even further, to approximately 52.6°C, by the introduction of either S162P and/or V323A in variant CbbM(1234). However, the second, higher melting point was slightly lowered in CbbM(1234) compared to all other variants. Hence, it is not clear if S162P and/or V323A might stabilize parts of the protein while destabilizing others.

Spectrophotometric *in vitro* assays testing for a change of kinetic enzyme parameters in an aerobic or anaerobic atmosphere did not show relevant differences between tested variants besides a slightly reduced turnover for variant CbbM(1234) under aerobic conditions (Fig. 5D, Supp. Fig. S16 and S17). This might be due to a reduced specificity *S(℅)* of the respective variant. Hence, we assume that changes observed in the pooled growth in *Synechocysits* among the 16 different variants are mainly based on their altered stability and expression levels.

### Establishment of a photoautotrophic high-throughput Rubisco screening platform

The suggested amino acid exchanges and resulting combinatorial mutant variants gave some insight into the protein fitness landscape of *Gallionella* CbbM. Two of the exchanges suggested by EVmutation helped to rescue the detrimental effect of the other two exchanges and might also contribute more generally to an increased robustness against detrimental replacements. This shows that EVmutation coupled to “*in silico* evolution” may be a valuable aid in deciding on amino acid exchanges to be tested in an experimental set-up limited to a low number of positions or exchanges. Future studies should test the used screening system at further growth conditions. Different growth conditions, such as mixotrophy as well as short or prolonged phases of darkness, have a strong impact on *Synechocystis* metabolite levels (13–16). In prolonged darkness or in fluctuating light, it is likely that CbbM might not react to the *Synechocystis* host cell’s signals aimed at the (in)activation or (re)activation of Rubisco. Overly active Rubisco in dark phases might be detrimental due to increased 2PG accumulation and energy-intensive photorespiration (16). Screening at these conditions might reveal enzyme variants that are better integrated into the host’s metabolism.

Furthermore, our established screening system enabled us to test a pool of different CbbM variants in parallel *in vivo* in a photoautotrophic host. In a previous study, we successfully followed the growth of approximately 20,000 sgRNA variants in a pooled CRISPRi screen using a similar growth competition set-up (32). Hence, we are confident that screening larger Rubisco variant libraries is feasible using the presented platform to elucidate a larger proportion of the Rubisco protein landscape, which might enable to optimize cyanobacterial or even plant growth and productivity in the long run.

## Methods

### Prediction of beneficial higher-order mutant variants using EVmutation and *in silico* evolution

The amino acid sequence of *Gallionella* Rubisco, CbbM, (GenBank identifier OGS68397.1) (51) was used as input for the EVmutation (24, 25) web server (https://v2.evcouplings.org/) with default settings (bitscores 0.1, 0.3, 0.5, 0.7; 5 search iterations; sequence database UniRef90 (52); position filter 70%; sequence fragment filter 50%; removing similar sequences 90%; downweighting similar sequences 80%; statistical inference model: pseudo-likelihood maximization). The web server identified a bitscore of 0.5 as optimal for alignments, which led to a multiple sequence alignment of 1,905 sequences (N_eff_/L = 4.9). The created model and multiple sequence alignment were downloaded and the inferred fitness landscape was navigated using *in silico* evolution, i.e. a Metropolis-Hastings Monte Carlo Markov chain (27), to identify higher-order mutant variants with predicted fitness values higher than the wild-type sequence. We decided to use this algorithm to explore the fitness landscape given by the Potts model created by EVmutation, but it can also be combined with other protein fitness prediction models, e.g. DeepSequence. The used code is available on https://github.com/ute-hoffmann/EVmut_inSilico and Zenodo (doi: 10.5281/zenodo.13971225). Briefly, an *in silico* evolution trajectory consists of multiple mutation steps. The algorithm started with the wild-type protein sequence of CbbM as the initial state. In each following step, a random amino acid exchange was proposed and the fitness value of the resulting sequence was predicted by EVmutation. Otherwise, with a certain acceptance probability given by the temperature value, the step was accepted nonetheless. Hence, the algorithm can help to traverse valleys of protein sequences with low predicted fitness values and escape local fitness maxima. This would not be possible in the case of a greedy algorithm that only accepts proposed sequences with increased fitness values. A trust radius determined how many amino acids could be exchanged in total. To determine amino acid positions of interest, 2,000 trajectories were run with 500 steps each and a trust radius of two for three different temperatures (0.3, 0.03, 0.003). Based on these results, a temperature of 0.03 was used for further analyses. The sequences with the highest predicted fitness values were extracted from the trajectories and the introduced amino acid exchanges were identified and counted. The thirty most frequently occurring replacements were used for another run of *in silico* evolutions, in which possible amino acid exchanges were limited to these respective positions. In this second run, 750 trajectories were run with 500 steps and a trust radius of three. The amino acid exchanges which were co-occurring most frequently among the highest scoring sequences were picked for library construction.

For visualization of the amino acid exchanges of interest, we used the AlphaFold 2.0 structure prediction (53, 54) of CbbM (https://alphafold.ebi.ac.uk/entry/A0A1G0AW29) and the PyMOL Molecular Graphics System, Version 2.5.0 (Schrödinger, LLC.). Multiple sequence alignments were created using Clustal Omega provided by EMBL-EBI (55) and analyzed using Jalview (v2.11.0) (56). Figures were created using Inkscape (v1.1).

### Bacterial strains and general culture conditions

We used a non-motile glucose-tolerant wild type of *Synechocystis*, which was kindly provided by Pauli Kallio, University of Turku. For culturing, BG-11 medium (57) supplemented with HEPES buffer (20 mM, pH 7.8) was used. If not cultivated in a photobioreactor or indicated otherwise, cultures were grown at 30°C under continuous white-light (45 µmol photons m^−2^ s^−1^) in ambient air enriched with 1% (v/v) CO_2_. Liquid cultures were grown in flat base uncoated 6-well plates (Sarstedt) or Erlenmeyer flasks under constant shaking (160 rpm). Plate cultures were grown on 1% (w/v) bacto-agar BG-11 plates containing 0.3% (w/v) sodium thiosulfate. Kanamycin (50 µg ml^−1^), spectinomycin (30 µg ml^−1^), gentamicin (2 µg ml^−1^), erythromycin (20 µg ml^−1^) and anhydrotetracycline (aTc, concentrations as indicated) were added to plate and liquid cultures when needed. For construction of plasmids carrying the *SPdcas9* gene, the CopyCutter EPI400 (Biosearch Technologies (Lucigen)) *E. coli* cell line was used. For all other cloning, the XL1-Blue *E. coli* cell line was used. For protein expression, *E. coli* BL21(DE3) were used. When appropriate, kanamycin (50 µg ml^−1^) was added to *E. coli* cultivation media.

### Construction of mutant strains

All *Synechocystis* strains and relevant oligonucleotides used in this study are listed in supplementary Tables S18 and S19. Annotated sequences of plasmids used for strain engineering are provided in .gb file format as supplemental information. All pEEK2-based plasmids (58) were transferred into receiver *Synechocystis* strains by triparental mating with *E. coli* XL1-Blue harboring the constructed plasmids and *E. coli* HB101 helper cells with the plasmid pRL443-Amp^R^ as described in (58). All other plasmids were introduced into the respective receiver strains by natural transformation.

All cloning was done by a combination of PCR, restriction enzyme digestions, ligations and AQUA cloning (59). Plasmids used for the insertion of a partial or complete CRISPRi system were derived from the plasmid pMD19T_psbA1_PL22_dCas9_B0015_SpR (33), which introduces the gene encoding catalytically dead Cas9 (dCas9, mutations D10A and H840A) from *Streptococcus pyogenes*, *SPdcas9*, as part of the tetR_PL22_dCas9_SpR expression cassette into the *slr1181* locus (*psbA1*). The annotated DNA sequence of the re-sequenced plasmid is given in file dcas-empty.gb. For targeting *rbcL* using CRISPR inhibition, the two protospacers with the lowest fitness values under highlight, high-CO_2_ (HL HC) and lowlight, low-CO_2_ (LL LC) conditions in a recent CRISPRi screen (32) were chosen: rbcL|10 (5’-ACGCCCGCCTTAAACCCTGCTT-3’) and rbcL|47 (5’-TCGGGGGTATAGTAGGTC-3’). To create the full CRISPRi system, both were placed under the control of an aTc-inducible L22 promoter (60). The DNA sequence of the plasmid introducing dCas9 and these sgRNAs is given in file dcas9_2xsgrna.gb. Before using the strain carrying the complete CRISPRi construct for downstream strain construction, a homozygous insertion of the construct into the genome was checked by cPCR using a small amount of cell material as template, DreamTaq 2X Green PCR Master Mix (Thermo Scientific) and primer pairs P01/P02 as well as P03/P04 (Supp. Fig. S18).

*Gallionella* Rubisco (GenBank identifier OGS68397.1) (51) was codon-optimized for expression in *Synechocystis* and synthesized as gBlocks gene fragment (idt). All *Gallionella* Rubisco variants were expressed from pEEK2-based plasmids under the control of the constitutive trc promoter. Point mutations were introduced into the *Gallionella* Rubisco coding sequence by performing PCRs using Phusion DNA polymerase (Thermo Scientific) and primer pairs M01/M02, M03/M04, M05/M06, M07/M08 and M09/M10, which were designed using NEBaseExchanger (NEB). Subsequently, 1 µl PCR product was mixed with 1 µl T4 DNA ligase buffer (10x), 1µl T4 PNK (10 U), 1µl T4 DNA ligase (5 U), 1µl DpnI (10 U) (all Thermo Scientific) and 5 µl H2O, incubated for 1 hour at room temperature and transferred to *E. coli* XL1-Blue. Higher order mutant variants were obtained by running several rounds of site-directed mutagenesis on the isolated plasmids of respective lower order mutant variants. Mutant variants were tagged with individual 20 nucleotide long barcodes which had been ordered as DNA fragments from idt.

The split GFP system was developed to minimize the potentially negative impact of GFP-tagging on a protein of interest (61). To this end, sfGFP was split into two parts and optimized for optimal assembly. DNA fragments with matching overhangs and encoding either of the two parts, i.e. the 11th alpha helix or alpha helices 1 to 10 of GFP, were ordered from idt and used to tag N-Strep-tagged CbbM C-terminally with GFP11 and create a plasmid integrating the gene encoding spGFP1-10, *spGFP1-10*, at the locus of *slr0168* under the control of a rhamnose-inducible promoter. The DNA sequence of the plasmid introducing the gene encoding Strep- and GFP11-tagged CbbM can be found in the file peek2-cbbm-gfp11.gb. For introducing either the rhamnose-inducible spGFP1-10 construct or, as a control, only a chloramphenicol resistance cassette, plasmids NS-GFP10 and sp3-CmR were created. Annotated DNA sequences of both plasmids can be found in files ns-gfp10.gb and sp3-cmr.gb.

Genes *ndhD3* and *ndhD4* were deleted using natural transformation (ndhd3_deletion_plasmid.gb and ndhd4_deletion_plasmid.gb). Full segregation of the resulting strains was tested by PCR using primer pairs P05/P06 for *ndhD3* and P07/P08 for *ndhD4* (Supp. Fig. S18) on DNA extracted using the GeneJET Genomic DNA Purification Kit (Thermo Scientific) according to the manufacturer’s Gram-positive extraction protocol.

### Microscopy

Liquid cultures of *cbbM*-GFP_11_|GFP_1-10_ and the control strain *cbbM*-GFP_11_|noGFP were diluted to an OD_750nm_ of 0.4 and expression of GFP_1-10_ was induced by the addition of rhamnose (4 mg ml^−1^) in one replicate of each culture. When cultures reached an OD_750nm_ of approximately 1.7, 2 ml of each culture and condition were harvested by centrifugation (6,000 x g, 5 min, 20°C). Supernatant was carefully decanted and pellets were resuspended in approximately 50 µl BG-11. Of these, 5 µl were spotted on a 1.8% (w/v) agarose (in phosphate-buffered saline) plate. These spots were dried for 10 to 30 minutes in a laminar flow cabinet. Spots were excised using a scalpel and inverted onto an uncoated 35 mm diameter petri dish with 20 mm diameter No. 1.5 glass bottom (MatTek). Confocal imaging was performed using a Zeiss LSM780 confocal microscope equipped with a 40X/1.2 NA water immersion objective and a spectral detector. GFP fluorescence was excited using a 488 nm laser, with emission captured between 493 and 630 nm. Autofluorescence was excited with a 633 nm laser, and emission was detected from 638 to 755 nm. The pinhole size was set to 1 Airy unit for both channels to ensure optimal optical sectioning and to maximize the signal-to-noise ratio. Images were analyzed using (Fiji Is Just) ImageJ 1.54f (62).

### Cultivation in photobioreactors

If indicated, cultivations in 8-tube Multi-Cultivator MC-1000-OD bioreactors (Photon System Instruments, Drasov, CZ) were performed using the pycultivator-legacy package (https://gitlab.com/mmp-uva/pycultivator-legacy) (63) and as described (32) with the following modifications: Light intensity was kept at 300 µmol photons m^−2^ s^−1^. If run in turbidostat mode, cultures were diluted if the turbidity threshold of OD_720nm_=0.2 was exceeded for two measurements in a row. Cultures were grown from the beginning of the cultivation at the indicated gas mixture. For growth competition assays of pools of several strains, CRISPRi repression was induced by adding aTc (0.2 µg ml^−1^) to cultivation tubes and feeding bottles after pools had acclimated and stable cultivation with regular back-dilutions was reached. Addition of aTc marked generation 0. After 24 hours, the aTc concentration was raised to 1 µg ml^−1^. Growth data was analyzed using the ShinyMC web application (https://github.com/m-jahn/ShinyMC, v0.1.1). Growth rates were calculated with “OD correction” set to true. The average growth rate of replicates was used for calculation of doubling times, i.e. the duration of one generation *t*, using the formula 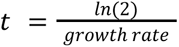. Cells were harvested after approximately the 2nd, 4th, 6th, 8th, and 10th generation by centrifuging 12 ml culture (4,500 x g, 4°C, 10 min). Supernatant was removed completely and cell pellets were frozen at −20°C until further usage.

For batch cultivations of strains *Syn*-sgRNA(−), *Syn*-sgRNA*_rbc_* and *Syn*-sgRNA*_rbc_ cbbM*(WT) at different gas conditions, pre-cultures were grown in ambient air enriched with 1% (v/v) CO_2_. After inoculation from plate cultures, liquid cultures were grown to an OD_730nm_ of approximately 0.8, diluted to an optical density OD_730nm_=0.2 and grown to OD_730nm_=0.8 to ensure a similar growth state of all used cultures. Subsequently, cultures were diluted to 0.2 and aTc (0.5 µg ml^−1^) was added. When cultures reached OD_730nm_=0.8, they were diluted to 0.1, aTc (0.5 µg ml^−1^) was added, and transferred into the bioreactors at the indicated gas feed conditions. Additional aTc (0.25 µg ml^−1^) was added every second day. For mass spectrometric analyses, 12 ml of culture was collected by centrifugation (4,500 x g, 4°C, 10 min) when an optical density of OD_720nm_=0.8 was reached.

For batch cultivations of strains *Syn*-sgRNA*_rbc_ cbbM*(WT), *Syn*-sgRNA*_rbc_ cbbM*(12), *Syn*-sgRNA*_rbc_ cbbM*(14), wild-type *Synechocystis* and Δ*ndhD3* Δ*ndhD4* strains, pre-cultures were grown at a gas feed of 5% (v/v) CO_2_, 75% (v/v) N_2_, 20% (v/v) O_2_ and a light intensity of 50 µE. Liquid cultures were diluted to OD_730nm_=0.4 and aTc added (1 µg ml^−1^). After 48 h, this was repeated. After another 48 h, cultures were diluted to OD_730nm_=0.05 and transferred to photobioreactors at the indicated gas feeds. Growth rates of batch cultures in photobioreactors was determined by performing a linear regression between 12.5 hours and 37.5 hours after transferral of cultures to the bioreactors using raw OD values and batch time as y and x values, respectively. One replicate of *Syn*-sgRNA*_rbc_ cbbM*(WT) was excluded from analyses at 1% (v/v) CO_2_, 78% (v/v) N_2_, 21% (v/v) O_2_, since its gas feed had stopped. Same holds for all replicates of wild-type Synechocystis at a gas feed of 1% (v/v) CO_2_, 99% (v/v) N_2_ and 0% (v/v) O_2_.

### Library preparation and next-generation sequencing

The GeneJET Plasmid-Miniprep-Kit (Thermo Scientific) was used to extract plasmids from cell pellets collected from turbidostat cultivations. Cell pellets were resuspended in 250 µl resuspension buffer and mixed with 50 µl of glass beads (diameter 425-600 µm, Sigma-Aldrich) in 1.5 ml protein LB SC Microtubes (Sarstedt). Samples were homogenized using a FastPrep-24 5G bead beating grinder and lysis system (MP Biomedicals) in three cycles of 45 s at 6.5 m s^−1^ with 30 s on ice between cycles. The plasmid extraction was continued according to the manufacturer’s protocol. DNA was eluted with 20 µl deionised 55°C-warm water. To amplify the barcode region of the plasmids by PCR, 9 µl of DNA eluate were used as template in a reaction volume of 30 µl using NEBNext Ultra II Q5 Master Mix (NEB) and PAGE-purified oligonucleotides S1 and an equimolar mixture of S2, S3 and S4 as forward and reverse primer, respectively. Oligonucleotides S2, S3 and S4 only differed regarding the length of a short stretch of random nucleotides (N, NN, NNN). Mixing the three oligonucleotides enabled phasing and to increase diversity in the resulting sequencing library. The manufacturer’s protocol for NGS PCRs and six elongation cycles were used. The PCR products were purified using Agencourt AMPure XP (Beckman Coulter, Inc.) according to manufacturer’s protocols and eluting DNA using 20 µl deionised water. In a second PCR, indices for multiplexing were added with the same PCR conditions as described above and using different combinations of oligonucleotides from the single and dual index sets of NEBNext® Multiplex Oligos for Illumina® (NEB) as primers. The amount of PCR product was quantified using a Qubit dsDNA HS Assay Kit and a Qubit 4 Fluorometer (both Thermo Scientific). Equal amounts of each sample were pooled and PCR product of the expected size was purified by performing a gel extraction from a 1.8% (w/v) agarose gel (in Tris acetate EDTA buffer, stained with Serva Electrophoresis DNA Stain G (Serva Electrophoresis)) using the GeneJET Gel extraction kit (Thermo Scientific) according to manufacturer’s instructions, except that elution was performed using 12 µl deionised 55°C-warm water. The concentration of PCR product was quantified using a Qubit TM 4 Fluorometer (Thermo Scientific) as described above and used for sequencing on a NextSeq 2000 system (Illumina) using NextSeq 1000/2000 P2 reagents v3 (Illumina) according to the manufacturer’s instructions.

### Analysis of sequencing data

Sequencing data was analyzed as described (32) by using the Nextflow pipeline (https://github.com/MPUSP/nf-core-crispriscreen commit e4aad5be10264d99e632761fc8fc56e68d6357c4 from 11th April 2024). Sequencing data was mapped to a library of the used barcodes. Compared to default settings, the error rate (parameter error_rate) was set to 0.2, the mapping quality cut-off (parameter filter_mapq) set to 1 and the sequence GTCTAGAatcgccgaaagtaattcaactccattaa…TCTAGATGCTTACTAGTTACCGCGGCCA was used for adapter trimming (parameter five_primer_adapter). Parameters run_mageck and gene_fitness were set to false. Further analyses were performed in the R programming language and are documented in R Markdown notebooks available at https://github.com/ute-hoffmann/CbbM_16variants (Zenodo doi: 10.5281/zenodo.14699592).

### Proteomics sample preparation

Cell pellets were collected as described for photobioreactor batch cultivations. Pellets were resuspended in 200 μl lysis buffer (100 mM HEPES pH 8, 1.5 M KCl, 3 mM MgCl) containing protease inhibitors (cOmplete, EDTA-free Protease Inhibitor Cocktail, Roche) and mixed with 100 μl glass beads (diameter 425-600 µm, Sigma-Aldrich). They were lysed by bead beating (FastPrep-24 5G lysis machine, MP Biomedicals) over six cycles of 45 s at 6.5 m s^−1^ and 4°C, with 30 s breaks on ice between cycles. The samples were then pelleted by centrifugation (21,000 x g, 5 minutes, 4°C) and supernatants were transferred to new tubes. Protein concentrations in the lysates were measured using a bradford assay (Bio-Rad protein assay dye reagent) using BSA as a standard at concentrations of 0, 0.05, 0.1, 0.2 and 0.4 mg ml^−1^. Lysates were diluted to a protein concentration of 2 mg ml^−1^ before being reduced with 18 mM DTT for 10 minutes at 96°C. Samples were then alkylated with 10 mM IAA for 30 minutes at 25°C before being diluted 4X in 100 mM ammonium bicarbonate. Subsequently, samples were digested by a trypsin/LysC protease mix (Pierce Trypsin/LysC protease mix, MS grade, Thermo Scientific) at a protein to protease mass ratio of 50:1 for 16 hours at 37 °C with 600 rpm of shaking, after which they were quenched with formic acid to pH < 2 and centrifuged (21,000 x g, 10 minutes, 25°C). Supernatants were then desalted through stage tips packed with six layers of C18 matrix using the following chromatographic workflow: activation with 50 μl acetonitrile, equilibration with 200 μl 0.1% formic acid, sample application, two times wash with 200 μl 0.1% formic acid and two times elution with 30 μl 80% acetonitrile, 0.1% formic acid. Samples were then evaporated to dryness at 40°C in a speedvac before being resuspended in 20 μl 0.1% formic acid and stored at −20°C until mass spectrometry analysis.

### Proteomics mass spectrometry analysis

Proteomics analysis was performed on a Q-exactive HF Hybrid Quadrupole-Orbitrap Mass Spectrometer coupled with an UltiMate 3000 RSLCnano System with an EASY-Spray ion source. 2 μL sample was loaded onto a C18 Acclaim PepMap 100 trap column (75 μm × 2 cm, 3 μm, 100 Å) with a flow rate of 7 μL per min, using 3% acetonitrile, 0.1% formic acid and 96.9% water as solvent. The samples were then separated on ES802 EASY-Spray PepMap RSLC C18 Column (75 μm × 25 cm, 2 μm, 100 Å) with a flow rate of 3.6 μL per minute for 40 min using a linear gradient from 1% to 32% with 95% acetonitrile, 0.1% formic acid and 4.9% water as secondary solvent. Mass spectrometry analysis was performed using one full scan (resolution 30,000 at 200 m/z, mass range 300–1200 m/z) followed by 30 MS2 DIA scans (resolution 30,000 at 200 m/z, mass range 350–1000 m/z) with an isolation window of 10 m/z. The maximum injection times for the MS1 and MS2 were 105 ms and 55 ms, respectively, and the automatic gain control was set to 3·106 and 1·106, respectively. Precursor ion fragmentation was performed with high-energy collision-induced dissociation at an NCE of 26 for all samples.

The prosit intensity prediction model “Prosit_2020_intensity_hcd” was used to generate a predicted peptide library from a FASTA file of the UniProt proteome set *Synechocystis* sp. PCC 6803: UP000001425, with the sequence of *Gallionella* CbbM added.

The raw spectra were converted to mzML using MSconvert and then searched using the EncyclopeDIA v. 1.2.2. search engine. Peptides detected in at least three replicates in every sample group were tested for differential peptide abundance using the MSstats package (version 4.12.0) in R (version 4.3.1.). For every peptide in each comparison MSstats estimated fold changes and p-values adjusted for multiple hypothesis testing (Benjamini-Hochberg method) with a significance threshold of 0.01.

During data analysis, one replicate taken at a gas feed of 1% (v/v) CO2, 78% (v/v) N2, 21% (v/v) O2 was removed from further analyses as an outlier.

### Western blot analysis

To ensure a similar growth status of different liquid cultures, cultures were grown twice to an OD_730nm_ of approximately 0.9 and subsequently diluted to an OD_730nm_ of 0.2. When cultures reached an OD_730nm_ of 0.9 for the second time, they were harvested by centrifugation (4,500 x g, 4°C, 10 min) and all remaining supernatant was removed. Cell pellets were frozen at −20°C until further usage. For lysis, pellets were resuspended in 200 µl phosphate-buffered saline containing protease inhibitors (cOmplete, EDTA-free Protease Inhibitor Cocktail, Roche) and mixed with 50 µl of glass beads (425–600 µm diameter, Sigma-Aldrich). Homogenization was performed as described for mass spectrometry analyses. Cell lysate was separated into soluble and insoluble fractions by centrifugation (20,000 x g, 4°C, 20 min). Protein concentrations were determined as described above using the Bio-Rad Protein Assay Dye Reagent Concentrate (Bio-Rad) and a BSA standard curve. Samples were analyzed using SDS-PAGE and western blot. Soluble fractions were mixed 1:1 with 2x Laemmli Sample Buffer (Bio-Rad) containing 50 mM dithiothreitol. After a 5 minutes 95°C denaturation step, approximately 20 µg protein of the soluble cell fraction was loaded onto precast 4–20% Mini-PROTEAN® TGX Stain-Free Protein Gels (Bio-rad) for SDS-PAGE. SeeBlue Plus2 prestained protein standard (Invitrogen) was used as size standard. Two identically prepared gels were run at the same time, of which one was stained using QC Colloidal Coomassie Stain (Bio-Rad) according to the manufacturer’s protocol. The second gel was used for western blotting onto a 0.2 µm PVDF membrane using Trans-Blot Turbo Mini 0.2 µm PVDF Transfer Packs (Bio-Rad) for 30 min at 25 V with an upper threshold of 1.0 A. After blocking in 5% milk powder in Tris-buffered saline containing Tween 20 (TBS-T, 20 mM Tris, pH7.5, 150 mM NaCl, 0.1% Tween 20), membranes were kept at 4°C overnight. Subsequently, they were shaken for 30 min at room temperature and washed twice with TBS-T for 10 minutes. Anti-strep-tag II rabbit IgG antibody (ab76949) in TBS-T (1:2000 dilution, 0.5 µg ml^−1^) was added to the membrane for 1 hour at room temperature. After two 10 minutes washing steps using TBS-T, secondary horseradish peroxidase (HRP) conjugated goat anti-rabbit IgG antibody (ab672) in TBS-T (dilution 1:5000, 0.16 µg ml^−1^) was added for 1 hour at room temperature. The membrane was washed again and Clarity Western enhanced chemiluminescence (ECL) substrates (Bio-Rad) were mixed 1:1 and applied to the membrane. The membranes were then analyzed with the Odyssey FC imaging system (Li-Cor).

### Protein expression and purification

The DNA sequence of the plasmid used for expression is provided online (pet28_rubisco.gb). *E. coli* BL21(DE3) were transformed with these pET28a-derived plasmids and for each CbbM variant, a single colony was used to inoculate 50 ml kanamycin-containing LB, which was incubated at 37°C overnight. The next day, 200 ml kanamycin-containing 2YT in a baffled flask were inoculated using preculture to an OD_600nm_ of 0.1 and kept shaking at 37°C. When reaching an OD_600nm_ of 0.6, 0.5 mM isopropyl-ß-D-thiogalactopyranosid (IPTG) was added to induce protein expression and the incubation temperature was reduced to 16°C. After 16 hours, cultures were harvested by centrifugation (3005 g, 4°C, 20 min). Cell pellets were resuspended in 10 ml B-PER^TM^ Complete Bacterial Protein Extraction (ThermoScientific) containing imidazole (20 mM) and incubated while rocking at room temperature for 25 min. The suspension was centrifuged (3005 g, 4°C, 20 min) and the supernatant was filtered using a 0.2 µm filter (SFCA +PF membrane, Corning). For each expressed variant, 2.5 ml Ni-NTA agarose beads (Qiagen) were prepared as described by the manufacturer. After three wash steps using B-PER^TM^ containing 20 mM imidazole, the filtered supernatant was added to the beads and incubated on ice for 1 h, while rocking. The bead suspension was transferred to a 5 ml polypropylene column (Qiagen) and the flow through was collected. The column was washed three times with 5x bead volume (6.25 ml) wash buffer (50 mM Tris-HCl, 20 mM imidazole, 500 mM NaCl, pH 7.4) and the wash flow through was collected. The column was washed once with 10% elution buffer and collected. The protein was eluted in four fractions of 0.5 ml elution buffer (50 mM Tris-HCl, 300 mM imidazole, 500 mM NaCl, pH 7.4) each. The fractions for each protein variant were collected and the buffer was exchanged to a Rubisco storage buffer (20 mM MgCl_2_, 20 mM EPPS, pH 8.0) using PD10 desalting columns (Cytiva) according to the manufacturer’s instructions. Eluates were concentrated using protein concentrators (10 kDa, Cytiva) by centrifugation (3005 g, 4°C) until reaching a volume of approximately 500 µl. Protein concentration was measured using the Bio-Rad Protein Assay Dye Reagent Concentrate (Bio-Rad) as described above and molarity calculated using molecular weights given in Supp. Table S20. Protein samples were used for downstream assays within 24 hours. Each protein purification was controlled by running SDS-PAGE as described above (compare Supp. Fig. S16 for an exemplary SDS gel).

### Nano differential scanning fluorimetry (nanoDSF)

The thermal stability of the Rubisco variants was determined by nano differential scanning fluorimetry (nanoDSF). The mutant variants were diluted to approximately 1 mg ml^−1^ with 100 mM EPPS and 50 mM NaHCO_3_. Three technical replicates of a blank sample (100 mM EPPS, 50 mM NaHCO_3_) and the mutant variants were transferred to Prometheus standard capillaries (NanoTemper Technologies) and placed in the Prometheus NT.48 instrument (NanoTemper Technologies). The thermal profile of the proteins was measured by the tryptophane shift and scattering at 350 nm and 330 nm in the temperature range from 20°C to 95°C at an increase of 1°C min^−1^ with 23% excitation power.

### Spectrophotometric Rubisco *in vitro* assay

Spectrophotometric *in vitro* assays were performed, with slight modifications, according to (10). Rubisco variant stocks were prepared at a protein concentration of 5 µM in 50 mM NaHCO_3_ and 100 mM EPPS. Assay components (see Supp. Table S21) were mixed (except RuBP and Rubisco protein) and 80 µl were added to each used well of a UV-transparent (Corning) 96 well plate. The absorbance at 340 nm was measured on a spectrophotometer (SpectraMaxR i3x, Molecular Devices or BioTek Epoch 2 microplate reader, Agilent) every 10 to 15 s for 15 minutes. To respective wells, 10 µl of the respective 5 µM Rubisco stock was added, mixed carefully by pipetting and absorbance at 340 nm was measured for 15 min. In a final step, 10 µl of RuBP was added to each well, all components were added by pipetting and absorbance was measured for 15 minutes. The assay was performed at 30°C in ambient air as well as in an anaerobic tent in an oxygen-free atmosphere (10 % CO2, 87.9 % N2, 2.1 % H2). A negative control containing RuBP without RubiscO and a positive control containing 1 mM 3PGA with Rubico were performed. The assay was performed trice for each condition. In the anaerobic tent, all tubes containing assay reagents (assay mix, protein, substrate) were opened and let to equilibrate for a minimum of 15 minutes before performing the assay.

The obtained absorbance values were converted with Lambert Beer (c = A/(ε*l)) with the extinction coefficient of 6.22 L mol^−1^ cm^−1^ for NADH and divided by two to account for two molecules of 3PGA produced per RuBP. In GraphPad Prism, linear regression was performed on concentration per second and the slope obtained was used to calculate the rate (*k_app_*= -(slope*10^9^)/protein concentration). As protein concentration, we used 500 nM.

## Abbreviations

2OG: 2-oxoglutarate
2PG: 2-phosphoglycolate
3PG: 3-phosphoglycerate
aTc: Anhydrotetracycline
CBB: Calvin-Benson-Bassham
CRISPRi: CRISPR interference
DMS: Deep mutational scan
*E. coli*: *Escherichia coli*
GO: Gene ontology
IPTG: Isopropyl-ß-D-thiogalactopyranosid
*k_C_*: Specificity constant of Rubisco for CO_2_
*K_C_*: Michaelis constant of Rubisco for CO_2_
*k_cat,C_*: Turnover number or catalytic constant of Rubisco for carboxylation reaction
*k_cat,O_*: Turnover number or catalytic constant of Rubisco for oxygenation reaction
*k_O_*: Specificity constant of Rubisco for O_2_
*K_O_*: Michaelis constant of Rubisco for O_2_
PCA: Principal component analysis
RbcLS: *Synechocystis* Rubisco
*R. rubrum*: *Rhodospirillum rubrum*
RuBP: Ribulose-1,5-bisphosphate
sgRNA: Single guide RNA
S(℅): Specificity factor of Rubisco for CO_2_ over O_2_
*Synechocystis*: *Synechocystis* sp. PCC 6803
TBS-T: Tris-buffered saline containing Tween20
*V_C_*: Limiting rate of Rubisco for CO_2_
*V_O_*: Limiting rate of Rubisco for O_2_

## Data Availability

Raw sequencing data were deposited at the European Nucleotide Archive (ENA accession number PRJEB79580). The mass spectrometry proteomics data have been deposited to the ProteomeXchange Consortium via the PRIDE (64) partner repository with the dataset identifier PXD059631 (Reviewer access details: **Project accession:** PXD059631 **Token:** 5D4oc2cOEiWh).

Source code used for analyses and some raw data is available on GitHub (https://github.com/ute-hoffmann/CbbM_16variants and https://github.com/ute-hoffmann/EVmut_inSilico) and Zenodo (dois 10.5281/zenodo.14699592 and 10.5281/zenodo.13971225).

## Supporting Information

Supporting Information includes:

- Supplementary Figures, including Figures supporting mass spectrometric findings, growth data and microscopy; and supplementary Tables including more details for mass spectrometric analyses, amino acid exchange effect prediction and strain construction (PDF)
- Supplementary Tables including the complete mass spectrometric data set, GSEA thereof, EVmutation predictions for CbbM and output of the sequencing analysis pipeline (.xlsx)
- Genbank (.gb) files of plasmids used for strain engineering

- dcas-empty.gb
- dcas9_2xsgrna.gb
- peek2-cbbm-barcode.gb
- pet28_rubisco.gb
- ndhd3_deletion_plasmid.gb
- ndhd4_deletion_plasmid.gb
- peek2-cbbm-gfp11.gb
- ns-gfp10.gb
- sp3-cmr.gb

## Author contributions

**UAH:** Conceptualization, Methodology, Software, Validation, Formal analysis, Investigation, Data Curation, Writing - Original Draft, Writing - Review & Editing, Visualization, Supervision, **AZS:** Methodology, Validation, Formal analysis, Investigation, Writing - Original Draft, Writing - Review & Editing, Visualization, **AK:** Methodology, Validation, Investigation, Writing - Review & Editing, **ES:** Methodology, Formal analysis, Investigation, Data Curation, Writing - Original Draft, Writing - Review & Editing, **HB:** Investigation, Writing - Review & Editing, **EE:** Investigation, Writing - Review & Editing, Funding acquisition, **POS:** Methodology, Writing - Review & Editing, Supervision, Funding acquisition, **EPH:** Conceptualization, Methodology, Writing - Original Draft, Writing - Review & Editing, Supervision, Funding acquisition

## Acknowledgements

We are grateful to Valentina Guccini for assistance with microscopy, Nick Crang for assistance with Illumina sequencing, Gustav Sjöberg for assistance with Rubisco *in vitro* assays, Michael Jahn for assistance with sequencing data analysis, Johannes Asplund for providing some of the code used in this manuscript, Rui Miao, Martin Hagemann, Annegret Wilde, Elisabeth Lichtenberg, Karen Schriever and Martha Falk for scientific discussions, Elsa Hesslow and Michele Russo for technical assistance and Elsa Hesslow for feedback on the manuscript.

