## Supplemental Info for "A cyanobacterial screening platform for Rubisco mutant variants"

#### Supplemental Information

**Supplemental Results & Discussion**

**Supplemental Figures**

**Supplemental Tables**

**Supplemental References**

### Supplemental Results & Discussion

##### Estimations on $CO_2$ and $O_2$ concentrations in microenvironment of CbbM

In *Syn*-sgRNA(-), the endogenous Rubisco variant and the enzyme carbonic anhydrase are encapsulated in carboxysomes, which are proteinaceous microcompartments that help to enrich free $CO_2$ around the enzyme and are thought to lower the local $O_2$ concentration (1). Deletion of a central carboxysome subunit, CcmM, prevents the formation of carboxysomes and leads to a cytoplasmic localization of Rubisco (2). A carboxysome-less mutant of *Synechocystis*, *ΔccmM*, in which Rubisco was localized cytoplasmically and was therefore more exposed to the relative gas feed composition, grew at a 50% reduced growth rate compared to wild-type cells in a gas feed of 5% $CO_2$ and air (2). Contrasting to *ΔccmM*, the growth rate of *Syn*-sgRNA(-) was largely unaffected by the composition of the external gas feed. This implies that carboxysomes in *Syn*-sgRNA(-) enable the cell to maintain a quasi constant gas environment around the enzyme at $CO_2$ feed levels between 1% and 5% and almost regardless of the presence of oxygen in the gas feed.

When expressed in *Synechocystis*, the Form II Rubisco from *R. rubrum* was localized cytoplasmically and not in the carboxysome (3). We confirmed the cytoplasmic localization of the Type II Rubisco *Gallionella* CbbM by microscopy (Supp. Fig. S2). Hence, *Gallionella* CbbM lacks the protective environment of the carboxysome, similar to *Synechocystis* Rubisco in *ΔccmM*.

It is difficult to determine the cytoplasmic concentration of gaseous $CO_2$ and $O_2$ relative to the external gas feed, particularly because oxygen is produced by photosynthesis when cells are grown under photoautotrophic conditions. Furthermore, while dissolved $CO_2$ can permeate the cell membrane and enter the cytosol, a portion is hydrated to carbonate by the NDH-1 complex leading to cytoplasmic $CO_2$ concentrations which might be lower than in the culture media (1). Extracellular concentrations might help to roughly estimate the intracellular concentrations.

Angermayr et al., (4) recorded a dissolved $CO_2$ at 145 μM in BG-11 when the gas feed was 0.5% (v/v). According to Henry's law, a gas composition of 1% $CO_2$ or 5% $CO_2$ could be expected to result in a proportional increase in dissolved $CO_2$ concentration to 290 μM and 1.5 mM, respectively. There are some estimates on the amount of $O_2$ generated by *Synechocystis* photosynthesis. Angermayr et al (4) showed that *Synechocystis* cultivated in turbidostat ($OD_{730nm}$ = 2.5) with a gas feed of 0.5% $CO_2$ and 99.5% $N_2$ generated oxygen so that $dO_2$ was 75 μM; the low OD in our cultivations ($OD_{720nm}$ = 0.2) would be expected to accumulate significantly less $dO_2$ (estimate 10 μM). This is significantly lower than $dO_2$ from air-saturated water (430 μM) and far below the estimated $CO_2$ concentration at both 1% as well as 5% $CO_2$ in the gas feed.

The $K_{M,C}$ for $CO_2$ of CbbM is reported to be 276 uM (5). Therefore, in a condition with 5% $CO_2$ and 95% $N_2$, we expect the enzyme to be saturated and the carboxylation rate proportional to $V_{max}$, while at a concentration of 1% $CO_2$ the carboxylation rate would reflect both $V_{max}$ and $K_M$.

A gas feed of 5% $CO_2$ and 20% $O_2$ should lead to approximately 1.5 mM dissolved $CO_2$ compared to air-saturated water with a $dO_2$ of 430 μM. Given a specificity constant for *Gallionella* Rubisco of S(%) = 10.0 ± 0.1 (5), the oxygenation reaction might play a relevant role at this condition and might determine the strain's growth rate.

#### Supplemental Figures

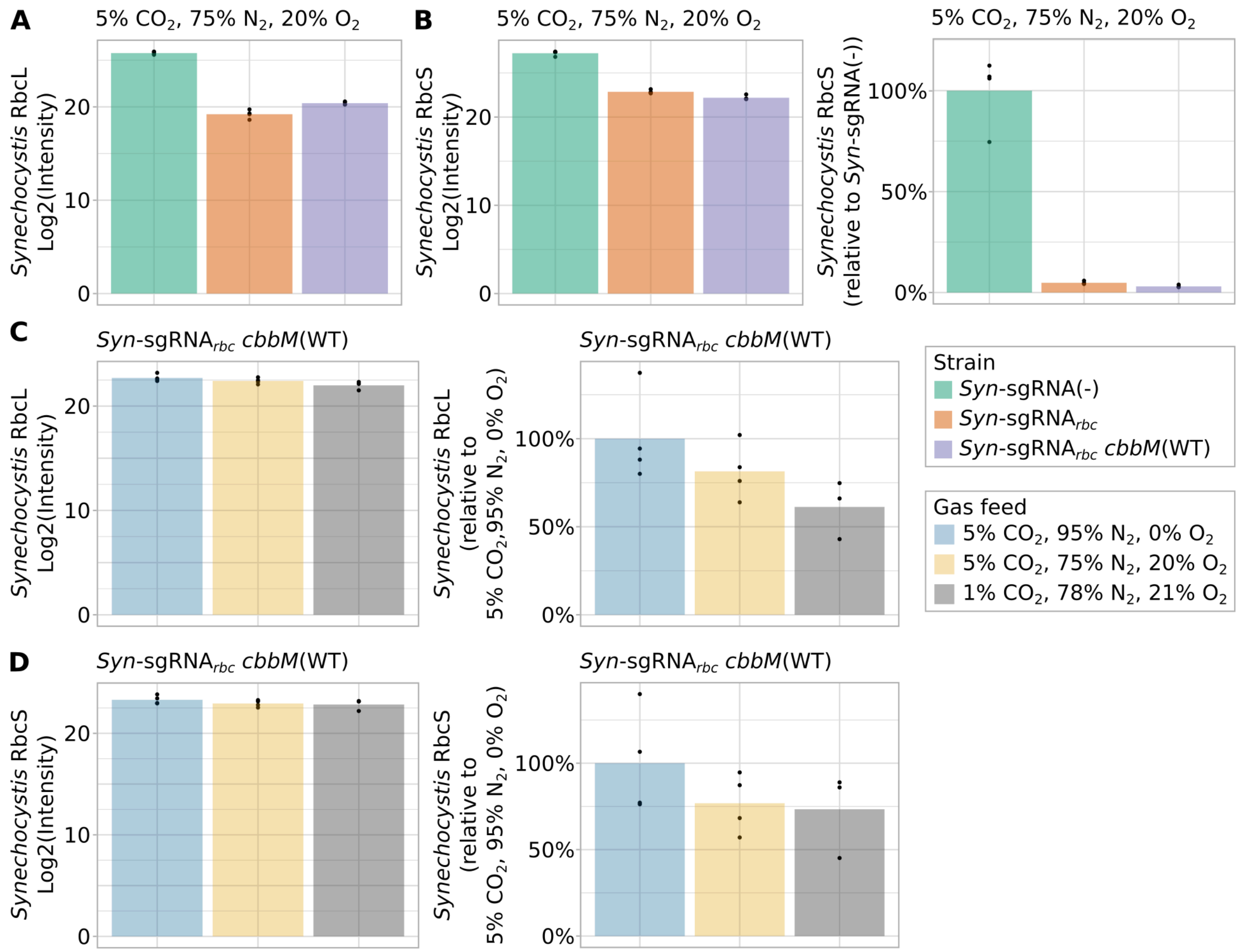


**Supp. Fig. S1 (related to Fig. 1B).** Mass spectrometry for *Synechocystis* Rubisco large and small subunits RbcL and RbcS. Panel (A), and left part of panels (B), (C), and (D): Log2 of measured signal intensities. Right part of panels (B), (C) and (D): relative amounts, normalized to *Syn*-sgRNA(-) (strain comparisons, panels A and B) or 5% $CO_2$, 95% $N_2$, 0% $O_2$ (gas feed comparisons, panels C and D). All cultivations were conducted at a light intensity of 300 µE. (**A**) Signal intensities for RbcL in different strains at 5% $CO_2$, 75% $N_2$, 20% $O_2$. (**B**) Data for RbcS in different strains at 5% $CO_2$, 75% $N_2$, 20% $O_2$. (**C**) Data for RbcL in *Syn*-sgRNA$_{rbc}$ *cbbM*(WT) at different gas feed compositions. (**D**) Data for RbcS in *Syn*-sgRNA$_{rbc}$ *cbbM*(WT) at different gas feed compositions.

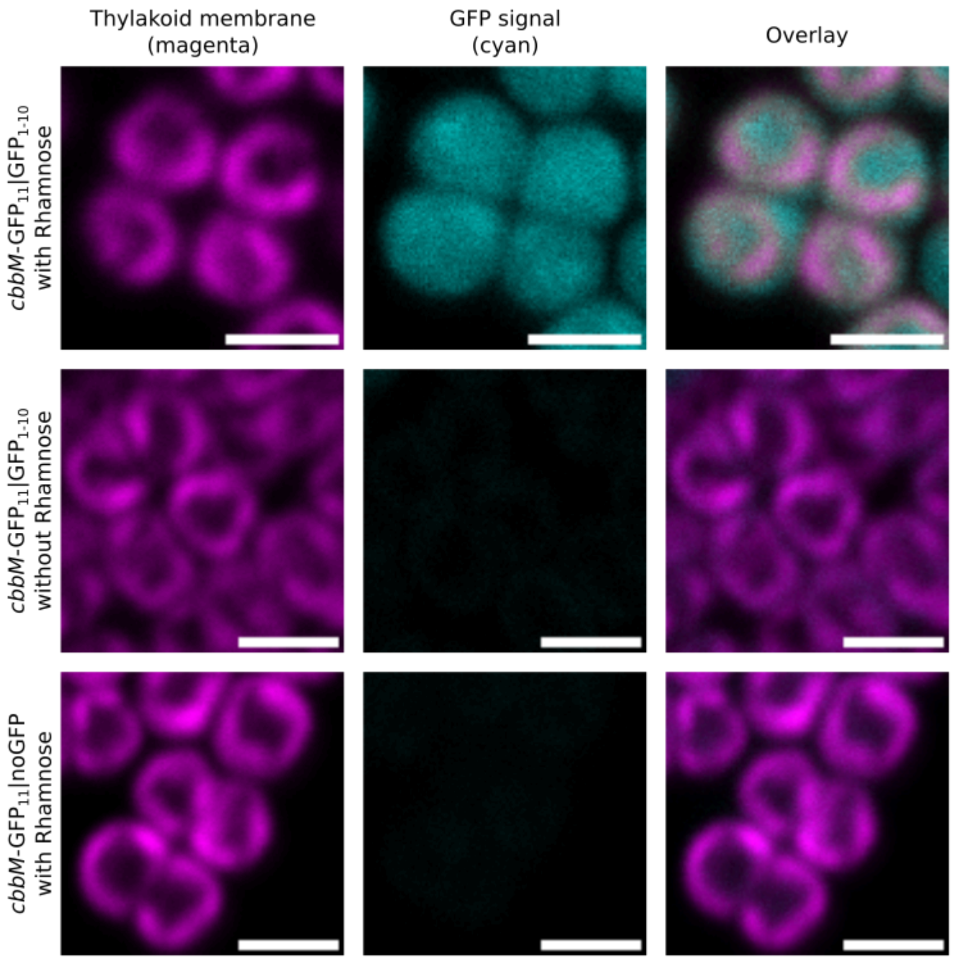


**Supp. Fig. S2.** Microscopy of strains expressing *Gallionella* CbbM fused to GFP11, *cbbM*-$GFP_{11}$|$GFP_{1\text{-}10}$ and *cbbM*-$GFP_{11}$|noGFP. In *cbbM*-$GFP_{11}$|$GFP_{1\text{-}10}$, $GFP_{1\text{-}10}$ expression is under the control of a rhamnose-inducible promoter ($P_{rha}$::$GFP_{1\text{-}10}$). In the control strain *cbbM*-$GFP_{11}$|noGFP, an antibiotic resistance cassette was introduced at the same genomic location instead of the $GFP_{1\text{-}10}$ construct. GFP fluorescence (cyan) and autofluorescence of the thylakoid membranes (magenta) is shown. All scale bars, 2 µm.

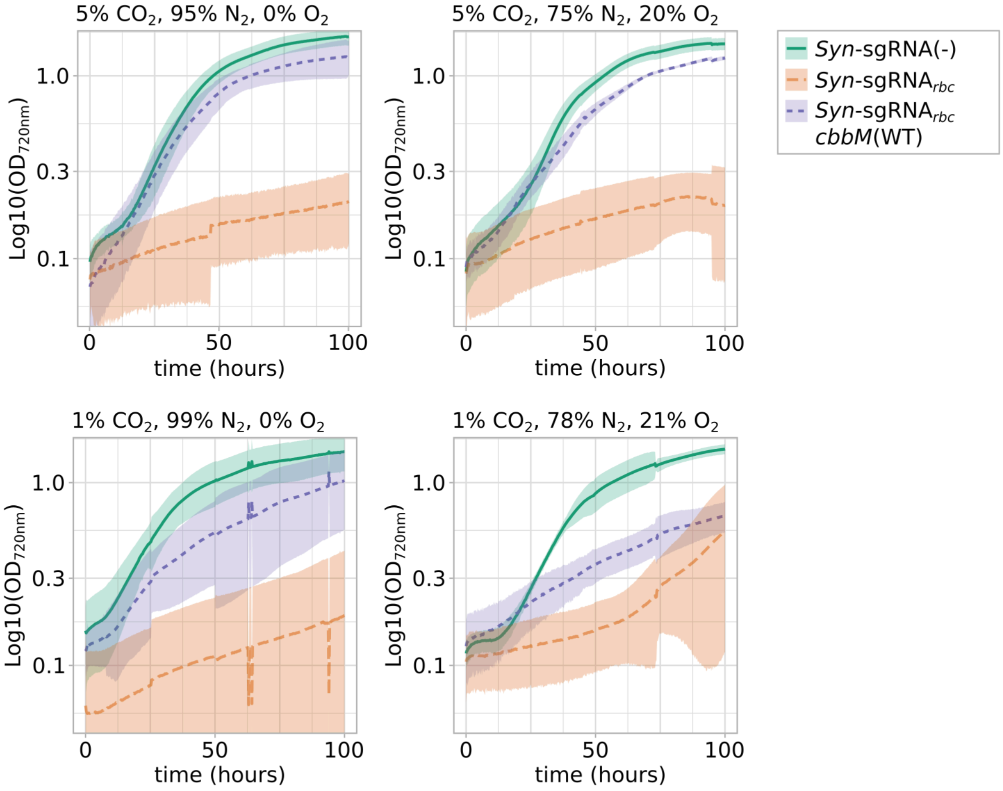


**Supp. Fig. S3 (related to Fig. 1C).** Growth curves of *Syn*-sgRNA(-), *Syn*-sgRNA$_{rbc}$ and *Syn*-sgRNA$_{rbc}$ *cbbM*(WT) at different gas feed $CO_2/O_2$ ratios after induction of the CRISPRi system using aTc (n=4, n=3 for *Syn*-sgRNA$_{rbc}$ *cbbM*(WT) at 1% $CO_2$, 78% $N_2$, 21% $O_2$). Growth is shown beginning from cultivating the strains at the indicated gas conditions. Light intensity was set to 300 µE. Shaded areas give the 95% confidence interval.

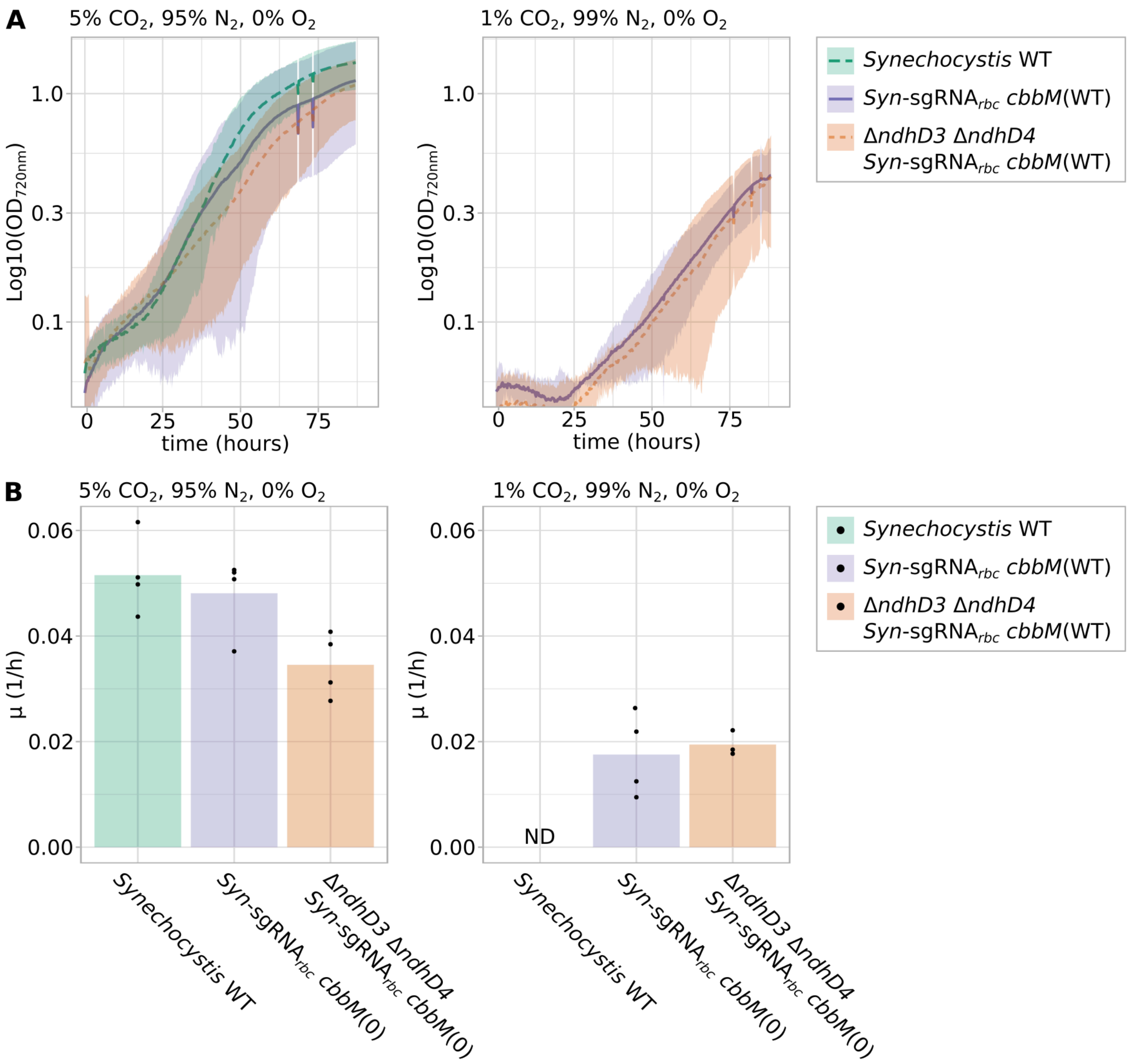


**Supp. Fig. S4.** Growth curves of wild-type *Synechocystis* (*Synechocystis* WT), *Syn*-sgRNA$_{rbc}$ *cbbM*(WT) and Δ*ndhD3* Δ*ndhD4* *Syn*-sgRNA$_{rbc}$ *cbbM*(WT) at different gas feed $CO_2/O_2$ ratios after induction of the CRISPRi system using aTc (n=4, n=3 for Δ*ndhD3* Δ*ndhD4* *Syn*-sgRNA$_{rbc}$ *cbbM*(WT) at 1% $CO_2$, 99% $N_2$, 0% $O_2$). Growth is shown beginning from cultivating the strains at the indicated gas conditions. Light intensity was set to 300 μE. Shaded areas give the 95% confidence interval. Growth of *Synechocystis* WT at 1% $CO_2$, 99% $N_2$, 0% $O_2$ was not determined (ND).

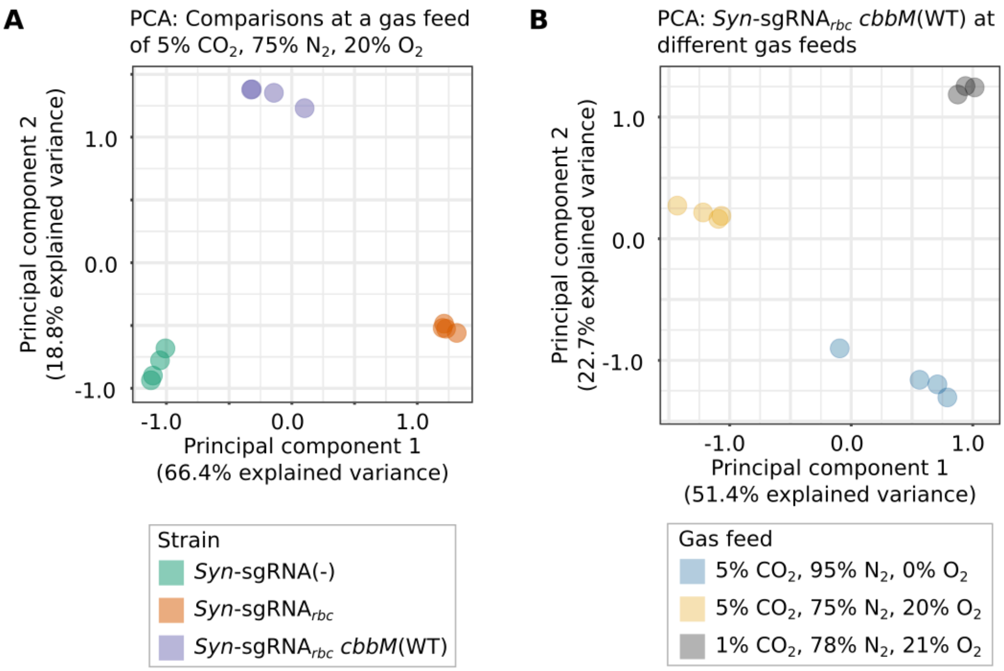


**Supp. Fig. S5 (related to Fig. 2).** Principal component analysis (PCA) of mass spectrometric data (Supp. Tables S1 to S4). (**A**) Comparison of different strains. (**B**) Comparison of *Syn*-sgRNA$_{rbc}$ *cbbM*(WT) at different gas feeds.

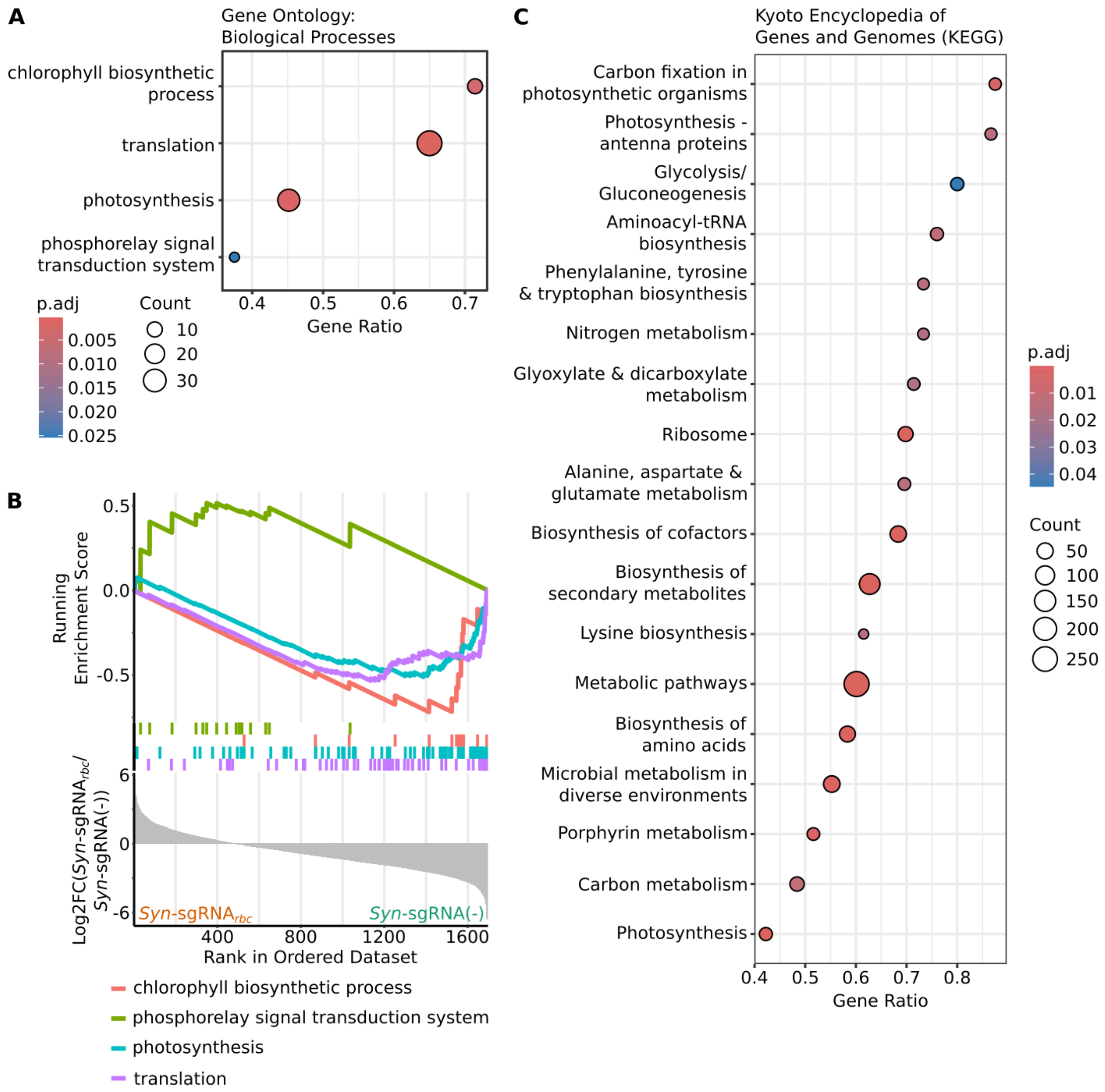


**Supp. Fig. S6 (related to Fig. 2).** Gene set enrichment analysis (GSEA) comparing mass spectrometric data of strains *Syn*-sgRNA(-) and *Syn*-sgRNA$_{rbc}$. As a metric to rank the detected proteins, Log2FC(*Syn*-sgRNA$_{rbc}$ / *Syn*-sgRNA(-)) was used. Terms which were significantly enriched in one of the two investigated strains are shown (*p.adj < 0.05*). (**A**) and (**B**) GSEA results for gene ontology (GO) terms describing biological processes. (**C**) GSEA results for Kyoto encyclopedia of genes and genomes (KEGG) pathways as annotated for *Synechocystis* sp. PCC 6803. All shown KEGG pathways were enriched in *Syn*-sgRNA(-) compared to

*Syn*-sgRNA$_{rbc}$. In panels (A) and (C), “gene ratio” describes the ratio of core enriched genes relative to the number of genes in a respective pathway, “count” describes the number of core enriched genes. In (B), the lowest part of the Figure depicts the whole set of detected proteins ranked by their Log2FC(*Syn*-sgRNA$_{rbc}$ / *Syn*-sgRNA(-)) values. The middle part of the figure shows color-coded where in the respective list the proteins belonging to the statistically significantly enriched GO term are located. The uppermost part shows the running GSEA enrichment score of these terms as a function of where in the ranked list the proteins are located. See also supp. Tables S5 and S6.

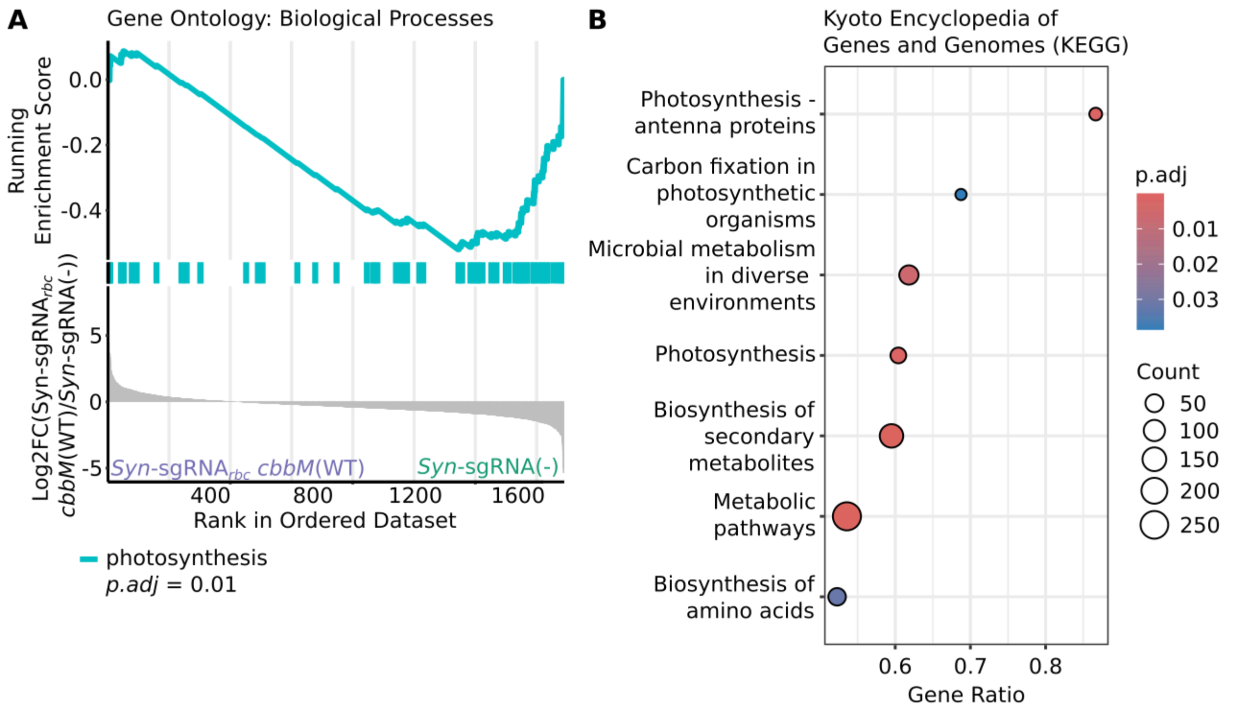


**Supp. Fig. S7 (related to Fig. 2).** Gene set enrichment analysis (GSEA) comparing mass spectrometric data of strains *Syn*-sgRNA(-) and *Syn*-sgRNA$_{rbc}$ *cbbM*(WT). As a metric to rank the detected proteins, Log2FC(*Syn*-sgRNA$_{rbc}$ *cbbM*(WT) / *Syn*-sgRNA(-)) was used. Terms which were significantly enriched in one of the two investigated data sets are shown (*p.adj* < *0.05*). (**A**) GSEA results for gene ontology (GO) terms describing biological processes. The lowest part of the Figure depicts the whole set of detected proteins ranked by their Log2FC(*Syn*-sgRNA$_{rbc}$ *cbbM*(WT) / *Syn*-sgRNA(-)) values. The middle part of the Figure shows color-coded where in the respective list the proteins belonging to the statistically significantly enriched GO term are located. The uppermost part shows the running GSEA enrichment score of these terms as a function of where in the ranked list the proteins are located. (**B**) GSEA results for Kyoto encyclopedia of genes and genomes (KEGG) pathways as annotated for *Synechocystis* sp. PCC 6803. All shown KEGG pathways were enriched for *Syn*-sgRNA(-) compared to *Syn*-sgRNA$_{rbc}$ *cbbM*(WT). “Gene ratio” describes the ratio of core enriched genes relative to the number of genes in a respective pathway, “count” describes the number of core enriched genes. See also supp. Tables S7 and S8.

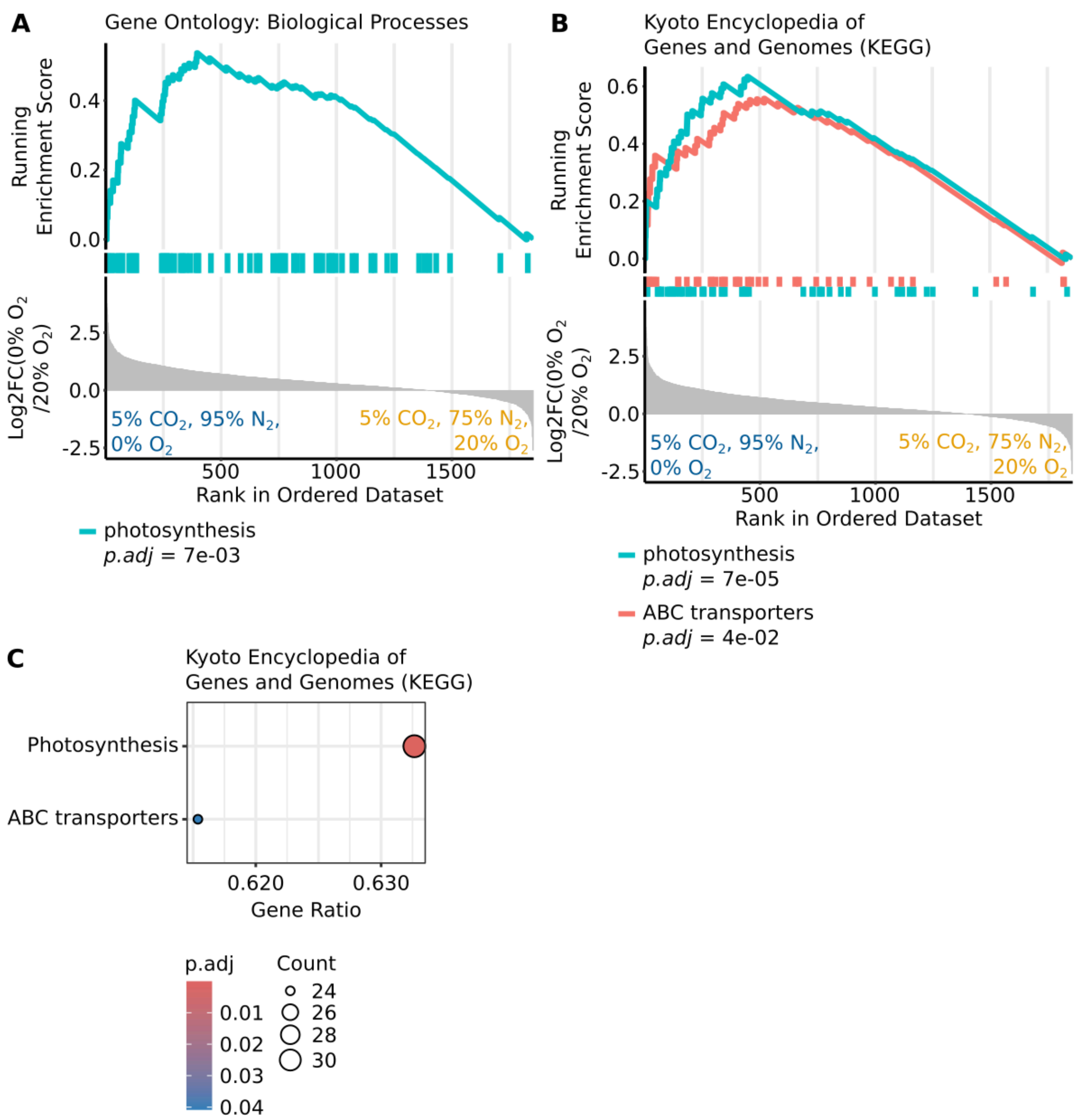


**Supp. Fig. S8 (related to Fig. 2).** Gene set enrichment analysis (GSEA) comparing mass spectrometric data of strain *Syn*-sgRNA$_{rbc}$ *cbbM*(WT) grown at a gas feed of either 5% $CO_2$, 95% $N_2$ and 0% $O_2$ or 5% $CO_2$, 75% $N_2$ and 20% $O_2$. As a metric to rank the detected proteins, Log2FC(5% $CO_2$, 95% $N_2$, 0% $O_2$ / 5% $CO_2$, 75% $N_2$, 20% $O_2$) was used. Terms which were significantly enriched in one of the two investigated strains are shown (*p.adj* < *0.05*). (**A**) GSEA results for gene ontology (GO) terms describing biological processes. (**B**) and (**C**) GSEA results for Kyoto encyclopedia of genes and genomes (KEGG) pathways. In (A) and (B), the lowest part of the figures depicts the whole set of detected proteins ranked by their Log2FC(5% $CO_2$, 95% $N_2$, 0% $O_2$ / 5% $CO_2$, 75% $N_2$, 20% $O_2$) values. The middle part of the figures shows color-coded where in the respective list the proteins belonging to the statistically significantly enriched terms are located. The uppermost part shows the running GSEA enrichment score of these terms as a function of where in the ranked list the proteins are located. In (C), “gene ratio” describes the ratio of core enriched genes relative to the number of genes in a respective pathway, “count” describes the number of core enriched genes. See also supp. Tables S11 and S12.

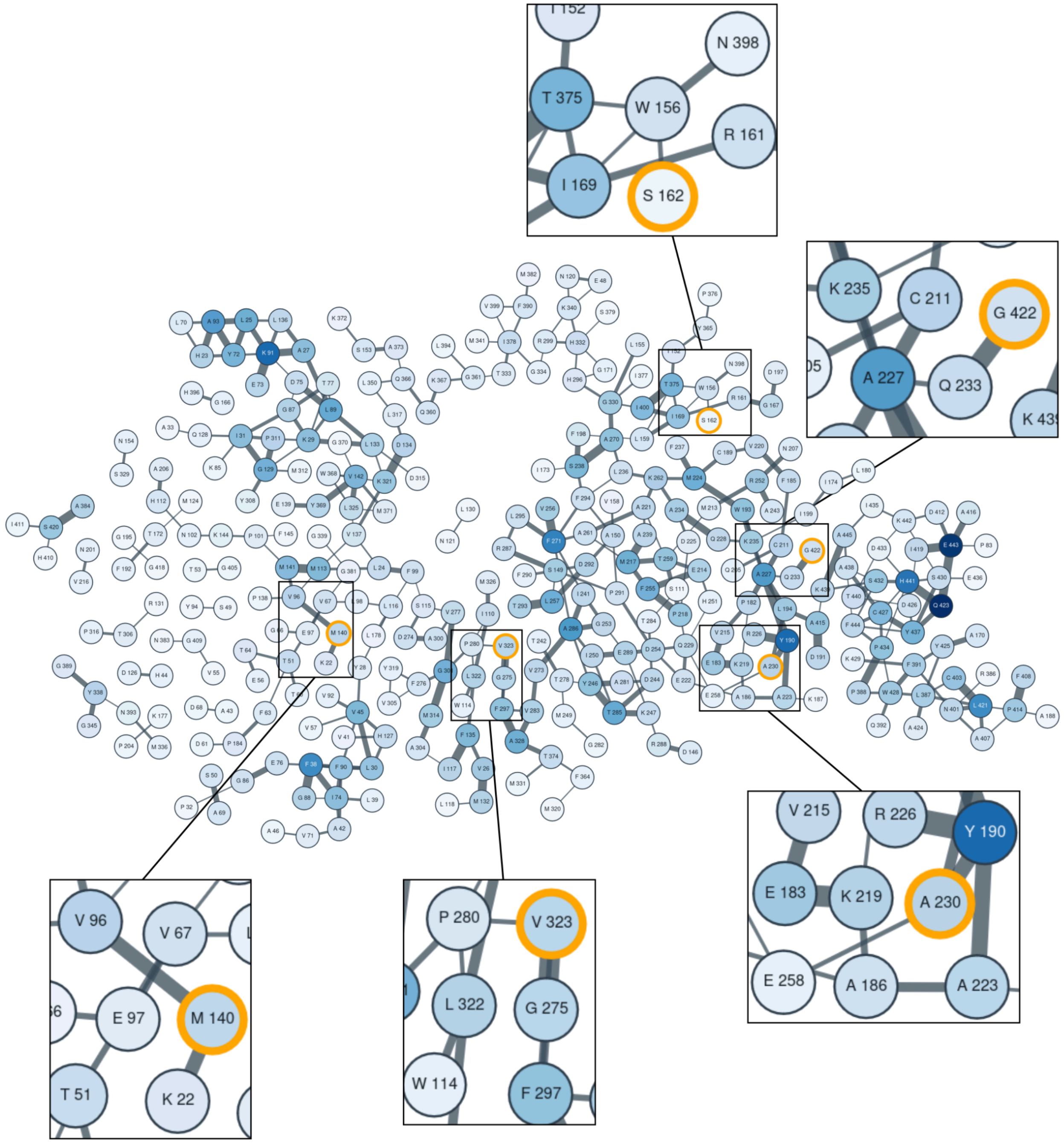


**Supp. Fig. S9 (related to Fig. 3).** Pair-wise couplings of amino acids in *Gallionella* Rubisco, CbbM, as given by the EVmutation web server (https://v2.evcouplings.org/). Amino acids investigated in this study are highlighted in orange. Blue color intensity and the thickness of edges indicate the degree of coupling to other residue positions. Inserts show a magnification of the immediate surroundings of the amino acid positions investigated in this study.

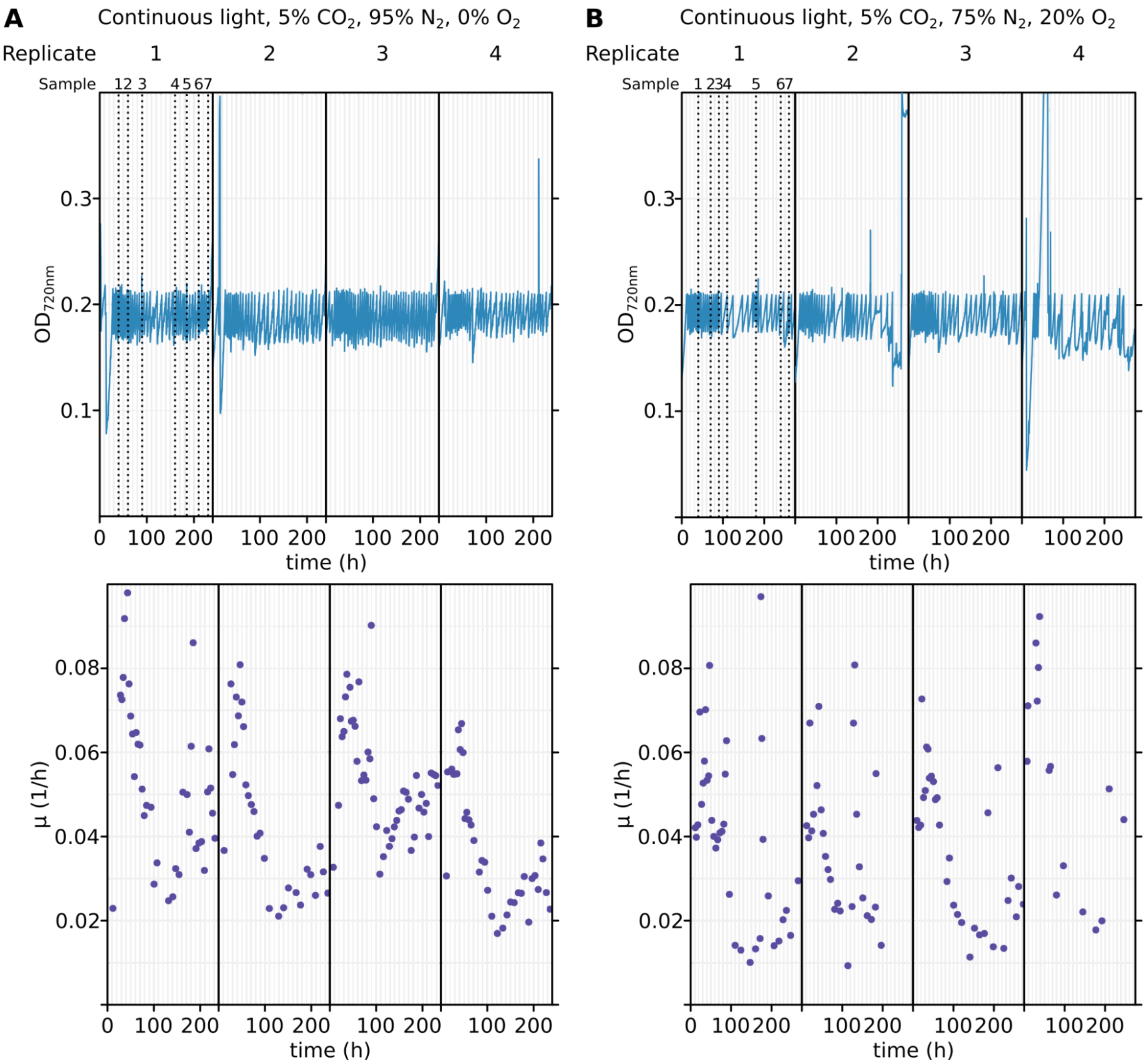


| Sample | Generation |
|---|---|
| 1 | 0 (+ aTc) |
| 2 | 2.2 |
| 3 | 4.1 |
| 4 | 5.2 |
| 5 | 7.3 |
| 6 | 8.8 |
| 7 | 10.0 |

| Sample | Generation |
|---|---|
| 1 | 0 (+ aTc) |
| 2 | 2.3 |
| 3 | 3.3 |
| 4 | 4.2 |
| 5 | 7.1 |
| 6 | 8.8 |
| 7 | 9.6 |

**Supp. Fig. S10 (related to Fig. 4).** Growth of mutant variant pool as recorded by ShinyMC. In both panels, optical density at 720 nm ($OD_{720nm}$) is given in the top part, and growth rates as calculated by ShinyMC in the bottom part. (**A**) Growth at continuous light of 300 μE, 5% $CO_2$, 95% $N_2$, 0% $O_2$. (**B**) Growth at continuous light of 300 μE, 5% $CO_2$, 75% $N_2$, 20% $O_2$.

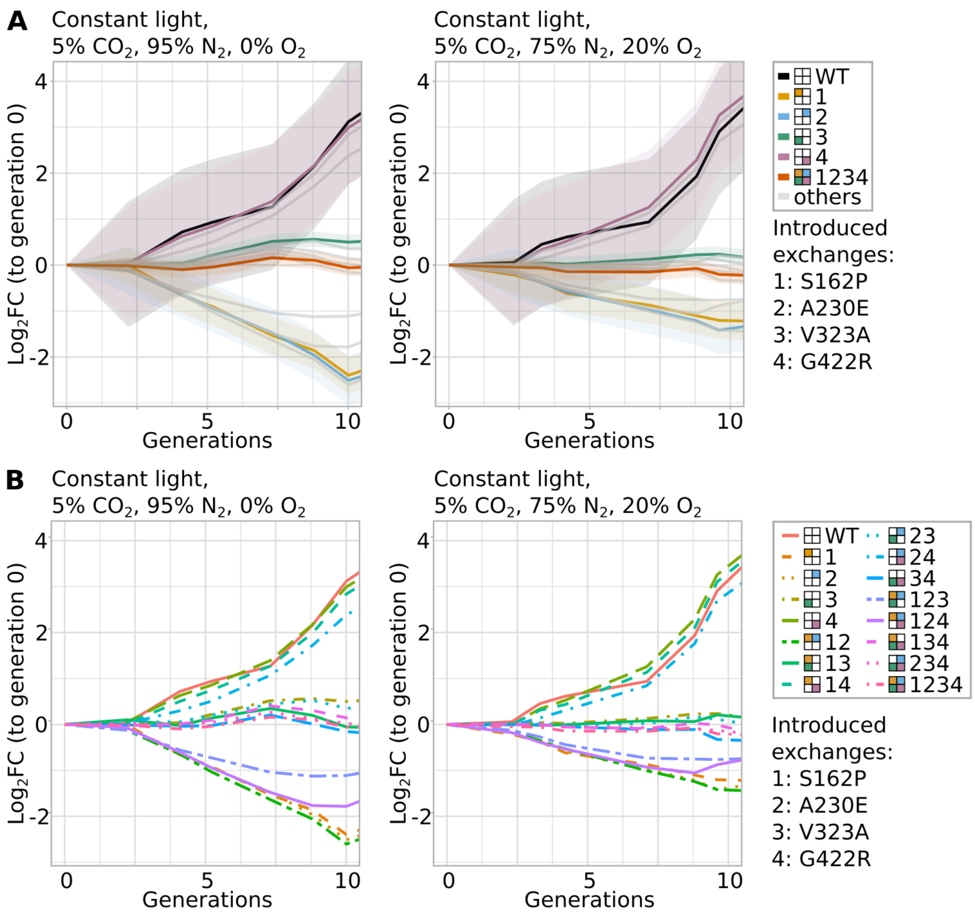


**Supp. Fig. S11 (related to Fig. 4).** Fitness values of competitive growth of the 16-member library at different growth conditions. All cultivations were performed at a light intensity of 300 µE. (**A**) Growth data including 95% confidence intervals as ribbons for wild-type CbbM, variants with a single amino acid exchange and of the quadruple mutant variant in color. Growth of other variants are plotted in light gray. (**B**) Fitness data of all variants present in the library. To make naming of higher-order mutants easier, different amino acid exchanges were assigned numbers based on their relative position in the protein's coding sequence and higher-order variants are called a combination of numbers: 1: S162P, 2: A230E, 3: V323A, 4: G422R. For instance, the variant containing S162P, V323A and G422R is referred to as 134.

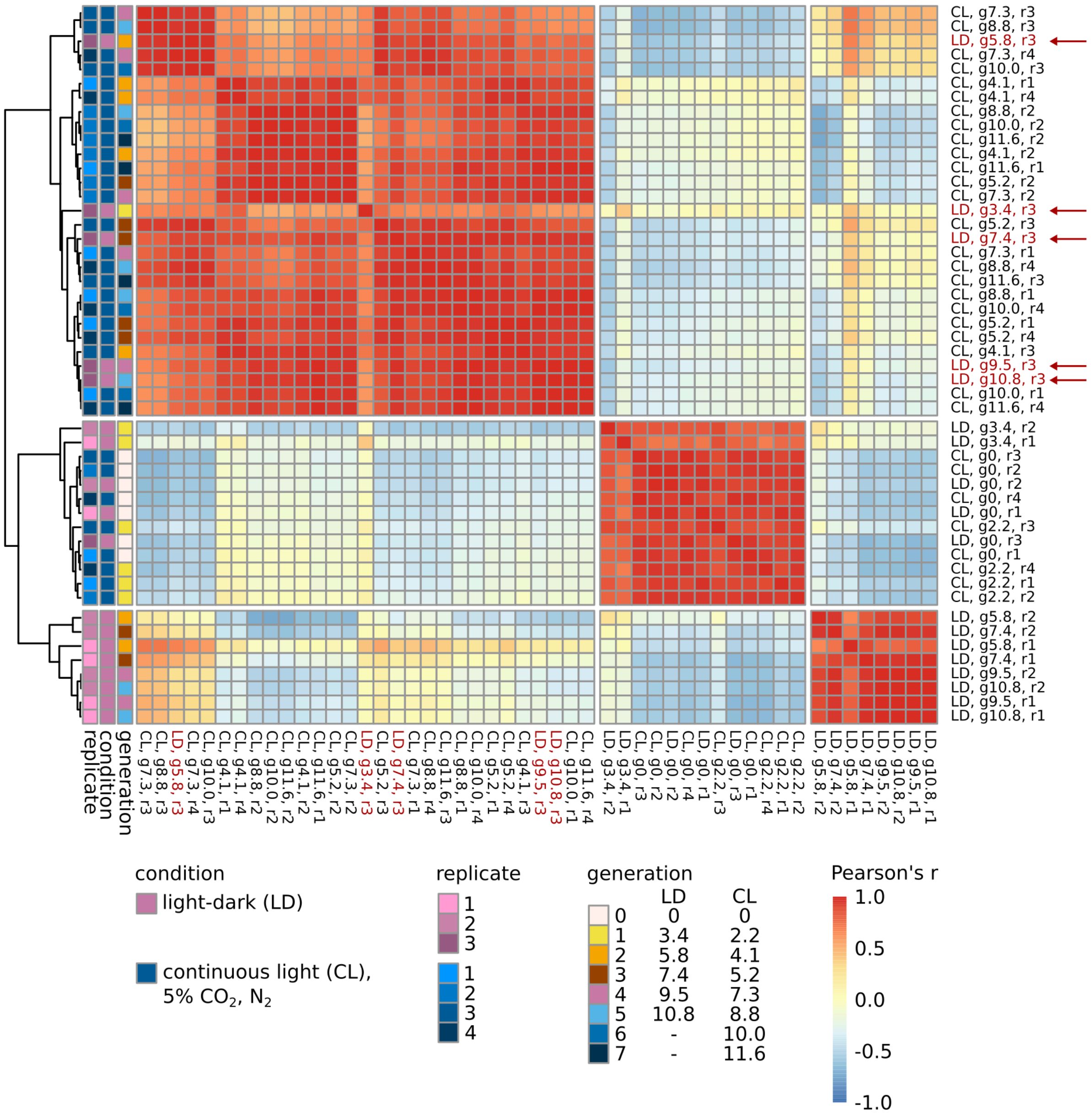


**Supp. Fig. S12 (related to Fig. 4).** Correlation between samples taken for cultivations with a gas feed of 5% $CO_2$, 95% $N_2$ and 0% $O_2$ at either continuous light (CL) or light-dark cycles (LD). Samples were clustered according to similarity. Samples for replicate number three for light-dark cycle cultivations showed a higher correlation with continuous light samples than with other light-dark replicates, probably due to stray light from the continuous light cultivation. Respective samples are indicated by red color and arrows. Since this reduced the number of usable light-dark samples to two, we decided to exclude this condition from downstream analyses.

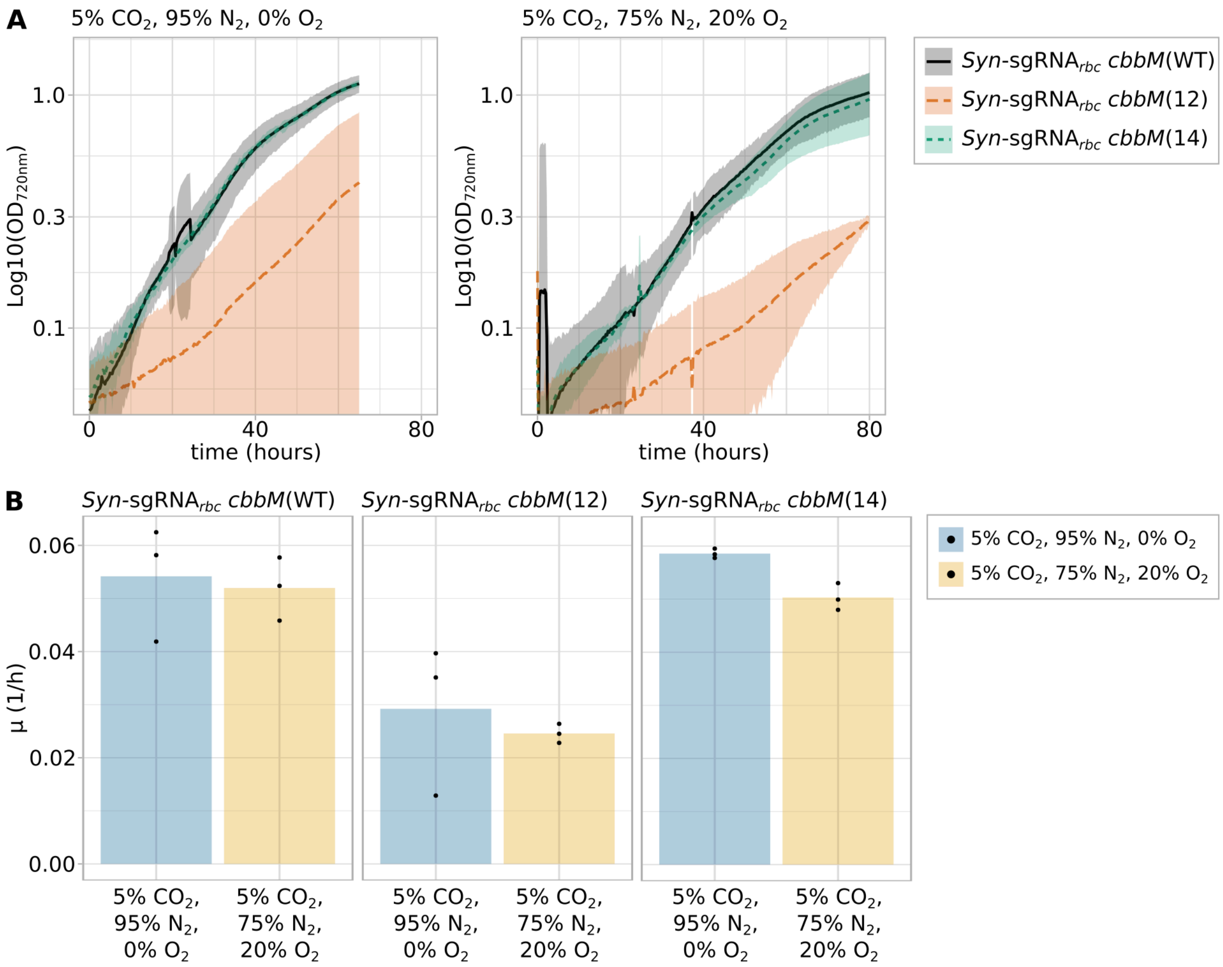


**Supp. Fig. S13.** Verification of pooled growth (**A**) Growth curves of *Syn*-sgRNA$_{rbc}$ *cbbM*(WT), *Syn*-sgRNA$_{rbc}$ *cbbM*(12) and *Syn*-sgRNA$_{rbc}$ *cbbM*(14) at a gas feed of 5% $CO_2$, 95% $N_2$, 0% $O_2$ (left) or 5% $CO_2$, 75% $N_2$, 20% $O_2$ (right) after induction of the CRISPRi system using aTc (n=3). Growth is shown beginning from cultivating the strains at the indicated gas conditions. Shaded areas give the 95% confidence interval. (**B**) Growth rates corresponding to growth curves shown in panel (A).

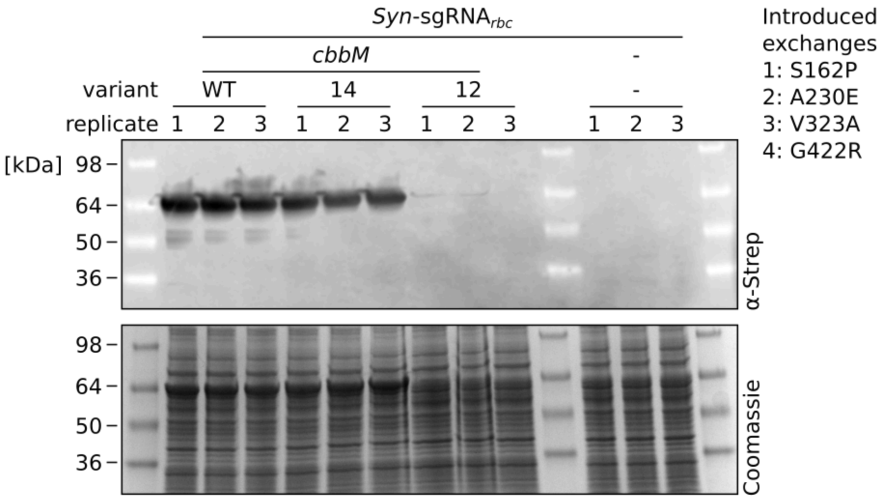


**Supp. Fig. S14 (related to Fig. 5).** Western blot analysis using horseradish peroxidase (HRP)-conjugated antibody directed against Strep-tag (αStrep) (top) and a Coomassie-stained SDS-PAGE gel (bottom) of the soluble cell fraction of strains expressing selected mutant variants to determine the mutant protein variants' relative abundance. Strain *Syn*-sgRNA$_{rbc}$ was used as a negative control. HRP chemiluminescence signal is overlaid with an image of the membrane. The protein of interest, N-Strep-tagged CbbM, has a size of 53 kDa. To make naming of higher-order mutants easier, different amino acid exchanges were assigned numbers based on their relative position in the protein's coding sequence and higher-order variants are called a combination of numbers: 1: S162P, 2: A230E, 3: V323A, 4: G422R. For instance, the variant containing S162P, V323A and G422R is referred to as 134.

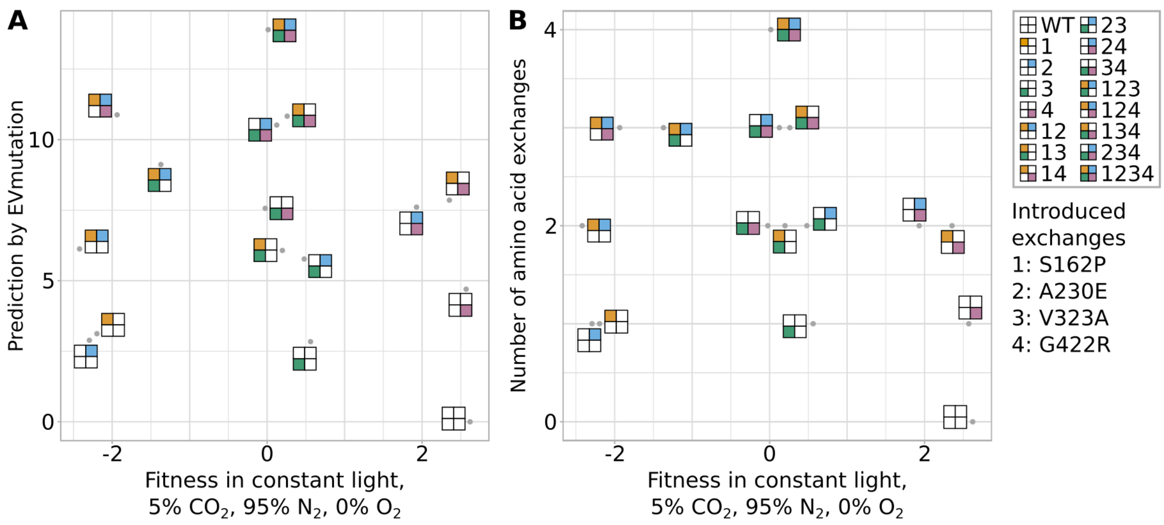


**Supp. Fig. S15.** (**A**) Comparison of EVmutation epistatic predictions and fitness values obtained for different variants by pooled growth at continuous light in a gas feed of 5% $CO_2$, 95% $N_2$ and 0% $O_2$ (compare Fig. 4). (**B**) Comparison of the number of introduced amino acid exchanges and fitness values obtained for different variants by pooled growth at continuous light in a gas feed of 5% $CO_2$, 95% $N_2$ and 0% $O_2$ (compare Fig. 4). For both panels, comparisons to the continuous light cultivation at 5% $CO_2$, 75% $N_2$, 20% $O_2$ are not shown since fitness values for the respective cultivations were highly correlated with the results for the cultivation using an oxygen-free gas feed. For reasons of clarity, we depict every variant as consisting of four boxes which represent the four different introduced amino acid exchanges. If a box is colored, the exchange was introduced and if left white, the exchange is absent. Furthermore, to make naming of higher-order mutants easier, different amino acid exchanges were assigned numbers based on their relative position in the protein's coding sequence and higher-order variants are called a combination of numbers: 1: S162P, 2: A230E, 3: V323A, 4: G422R. For instance, the variant containing S162P, V323A and G422R is referred to as 134.

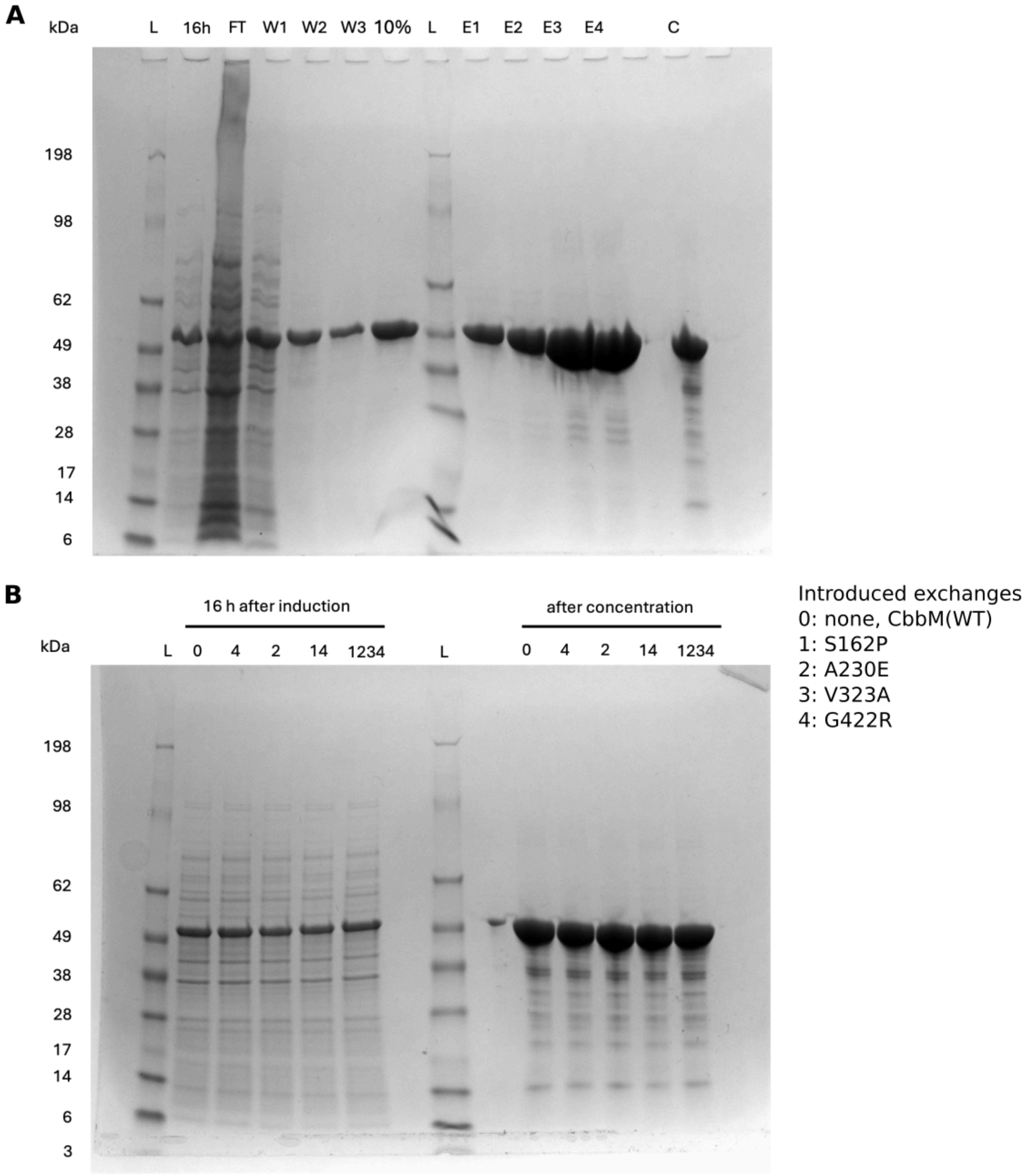


**Supp. Fig. S16 (related to Fig. 5C and 5D).** SDS-PAGE analysis of (**A**) one exemplary protein purification for CbbM(WT) and (**B**) protein lysates and concentrates of all purified protein variants. Gels were stained with Coomassie. Abbreviations are L: ladder, 16h: protein lysate 16 h post induction, FT: flow-through, W1 to W3: washing steps 1 to 3, 10%: washing step with 10% elution buffer, E1 to E4: eluate 1 to 4, C: concentrated eluate. Introduced amino acid exchanges are encoded as 0: none, 1: S162P, 2: A230E, 3: V323A, 4: G422R. For higher-order variants, these numbers were combined. For instance CbbM(14) combines exchanges S162P and G422R.

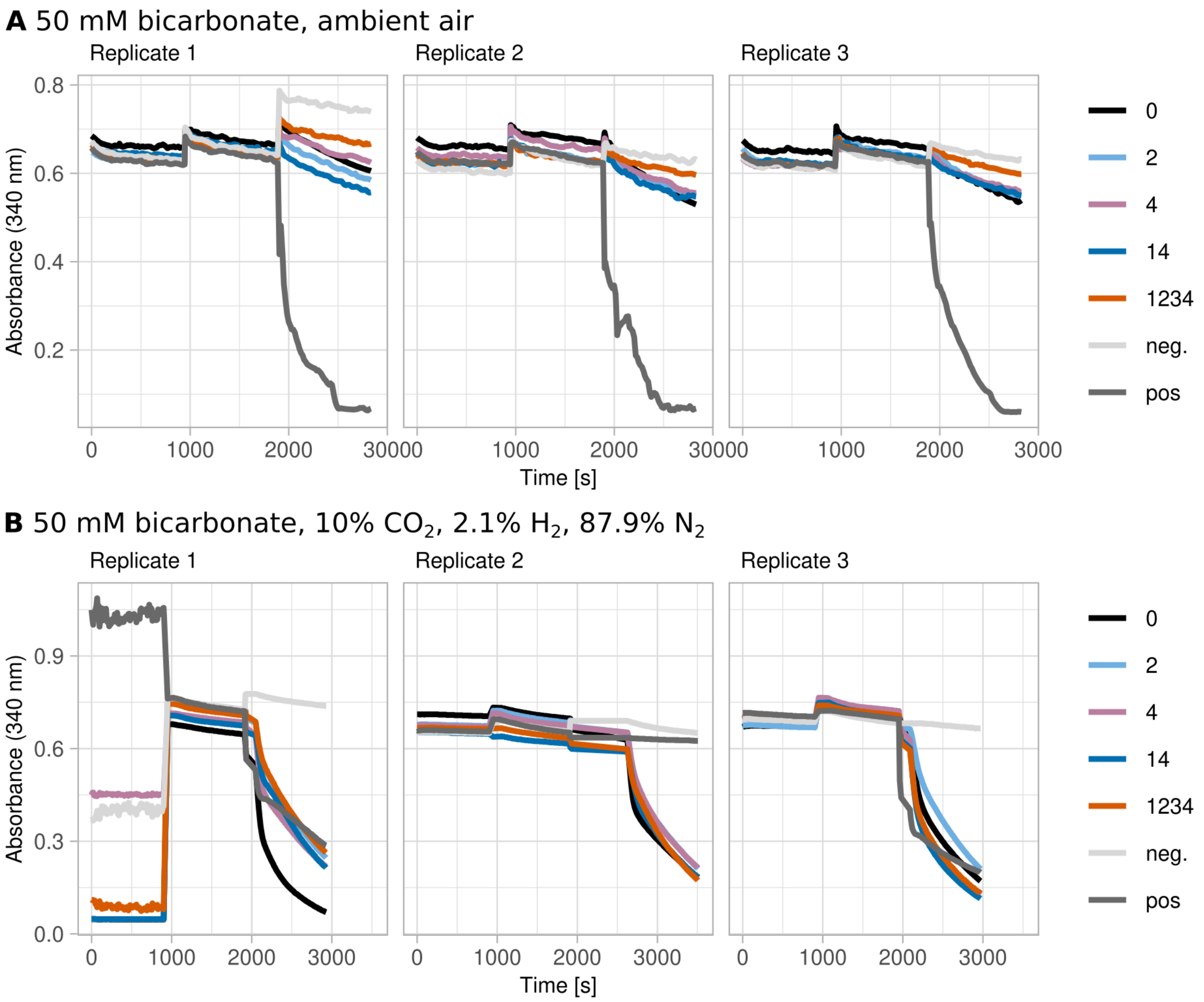


**Supp. Fig. S17 (related to Fig. 5D).** Spectroscopic data of the *in vitro* assays of the three measured replicates in (**A**) an oxygen-containing or (**B**) an oxygen-free atmosphere. The assays were conducted either in ambient air (panel A, approx. 0.04% $CO_2$, 78% $N_2$, 21% $O_2$) or in an oxygen-free gas atmosphere (panel B, 10% $CO_2$, 2.1% $H_2$, 87.9% $N_2$, 0% $O_2$). In both cases, 50 mM bicarbonate was added to the reaction mixture.

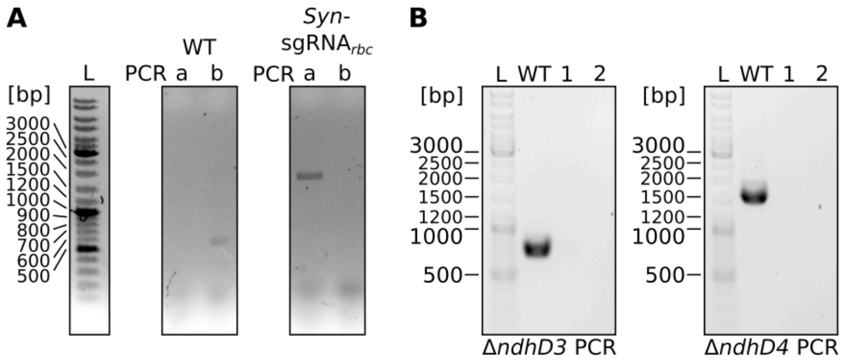


**Supp. Fig. S18.** Agarose TAE gels with products of PCR performed with primers to verify full segregation of strains used for further strain engineering and experiments. (**A**) PCRs run with primer pairs a) P01/P02, which gives no PCR product when the wild-type locus is present and a product of 1.6 kilobase pairs upon integration of the CRISPRi construct, and b) P03/P04, which gives a product of 528 base pairs with the wild-type locus and no band upon integration of the CRISPRi construct, on wild-type Synechocystis (WT) and *Syn*-sgRNA$_{rbc}$. (**B**) PCRs run with primer pairs P05/P06 (Δ*ndhD3* PCR) and P07/P08 (Δ*ndhD4* PCR) on WT and two clones of Δ*ndhD3* Δ*ndhD4* *Syn*-sgRNA$_{rbc}$ (numbered 1 and 2). Both primer pair combinations give no product if the respective gene was deleted and products of 738 base pairs (Δ*ndhD3* PCR) and approximately 1.6 kilobase pairs (Δ*ndhD4* PCR) when using the wild-type locus.

#### Supplemental Tables

For supplemental Tables S1 to 8, S11 to S14 and S16, see separate Excel workbook SuppTables_Synechocystis_CbbM-screening.xlsx. Tables are presented as an Excel workbook since they would exceed the limitation of a pdf file.

Supplementary Tables as part of separate Excel file:

- Supp. Table S1: Comparison of strains *Syn*-sgRNA(-) and *Syn*-sgRNA$_{rbc}$ at a gas feed of 5% $CO_2$, 75% $N_2$, 20% $O_2$ using mass spectrometry. Higher log2FC values indicate that a protein is more abundant in *Syn*-sgRNA$_{rbc}$. Column “Entry” gives the UniProtKB primary accession number. Columns “Locus tags”, “Gene names” and “Protein names” were computationally added based on these accession numbers and are not necessarily complete.
- Supp. Table S2: Comparison of strains *Syn*-sgRNA(-) and *Syn*-sgRNA$_{rbc}$ *cbbM*(WT) at a gas feed of 5% $CO_2$, 75% $N_2$, 20% $O_2$ using mass spectrometry. Higher log2FC values indicate that a protein is more abundant in *Syn*-sgRNA$_{rbc}$ *cbbM*(WT). Column “Entry” gives the UniProtKB primary accession number. Columns “Locus tags”, “Gene names” and “Protein names” were computationally added based on these accession numbers and are not necessarily complete.
- Supp. Table S3: Comparison of strain *Syn*-sgRNA$_{rbc}$ *cbbM*(WT) at gas feeds of 5% $CO_2$, 75% $N_2$, 20% $O_2$ and 5% $CO_2$, 95% $N_2$, 0% $O_2$ using mass spectrometry. Higher log2FC values indicate that a protein is more abundant in 5% $CO_2$, 95% $N_2$, 0% $O_2$. Column “Entry” gives the UniProtKB primary accession number. Columns “Locus tags”, “Gene names” and “Protein names” were computationally added based on these accession numbers and are not necessarily complete.
- Supp. Table S4: Comparison of strain *Syn*-sgRNA$_{rbc}$ *cbbM*(WT) at gas feeds of 5% $CO_2$, 75% $N_2$, 20% $O_2$ and 1% $CO_2$, 78% $N_2$, 21% $O_2$ using mass spectrometry. Higher log2FC

values indicate that a protein is more abundant in 1% $CO_2$, 78% $N_2$, 21% $O_2$. Column "Entry" gives the UniProtKB primary accession number. Columns "Locus tags", "Gene names" and "Protein names" were computationally added based on these accession numbers and are not necessarily complete.

- Supp. Table S5: Gene set enrichment analysis (GSEA) on the basis of gene ontology (GO) terms comparing *Syn*-sgRNA$_{rbc}$ and Syn-sgRNA(-). Higher values mean a term is enriched in *Syn*-sgRNA$_{rbc}$.
- Supp. Table S6: Gene set enrichment analysis (GSEA) on the basis of KEGG terms comparing *Syn*-sgRNA$_{rbc}$ and Syn-sgRNA(-). Higher values mean a term is enriched in *Syn*-sgRNA$_{rbc}$.
- Supp. Table S7: Gene set enrichment analysis (GSEA) on the basis of gene ontology (GO) terms comparing *Syn*-sgRNA$_{rbc}$ *cbbM*(WT) and Syn-sgRNA(-). Higher values mean a term is enriched in *Syn*-sgRNA$_{rbc}$ *cbbM*(WT).
- Supp. Table S8: Gene set enrichment analysis (GSEA) on the basis of KEGG terms comparing *Syn*-sgRNA$_{rbc}$ *cbbM*(WT) and Syn-sgRNA(-). Higher values mean a term is enriched in *Syn*-sgRNA$_{rbc}$ *cbbM*(WT).

Supp. Table S9 and S10 are part of this pdf file, see below.

- Supp. Table S11: Gene set enrichment analysis (GSEA) on the basis of gene ontology (GO) terms comparing *Syn*-sgRNA$_{rbc}$ *cbbM*(WT) at gas feeds with and without $O_2$, 5% $CO_2$, 95% $N_2$, 0% $O_2$ and 5% $CO_2$, 75% $N_2$, 20% $O_2$. Higher values mean a term is enriched in the condition without added $O_2$ (5% $CO_2$, 95% $N_2$, 0% $O_2$).
- Supp. Table S12: Gene set enrichment analysis (GSEA) on the basis of KEGG terms comparing *Syn*-sgRNA$_{rbc}$ *cbbM*(WT) at gas feeds with and without $O_2$, 5% $CO_2$, 95% $N_2$,

0% $O_2$ and 5% $CO_2$, 75% $N_2$, 20% $O_2$. Higher values mean a term is enriched in the condition without added $O_2$ (5% $CO_2$, 95% $N_2$, 0% $O_2$).

- Supp. Table S13: Gene set enrichment analysis (GSEA) on the basis of KEGG terms comparing *Syn*-sgRNA$_{rbc}$ *cbbM*(WT) at gas feeds with and without $O_2$, 5% $CO_2$, 78% $N_2$, 21% $O_2$ and 5% $CO_2$, 75% $N_2$, 20% $O_2$. Higher values mean a term is enriched in the condition with a low $CO_2$ concentration (1% $CO_2$, 78% $N_2$, 21% $O_2$).
- Supp. Table S14: EVmutation predictions for wild-type CbbM, output from web server https://v2.evcouplings.org/ and downloaded on 13th May 2022.

Supp. Table S15 is part of this pdf file, see below.

- Supp. Table S16: Output of nf-core-crispripipeline giving the fitness values of different CbbM mutant variant strains at different conditions (HL_5N2: continuous light, gas feed 5% $CO_2$, 95% $N_2$, 0% $O_2$; HL_5O2: continuous light, gas feed 5% $CO_2$, 75% $N_2$, 20% $O_2$, pool: pool of all strains before inoculation of photobioreactors). For different time points (measured in generations), DESeq2 output is given (baseMean, log2FoldChange, lfcSE, stat, pvalue, padj).

Supp. Tables S17, S18, S19, S20 and S21 are part of this pdf file, see below.

**Supp. Table S9:** Subset of Supplementary Tables S3 and S4 with data for the Rubisco operon, carboxysome, carbonic anhydrase, CmpR and NdhR regulon according to (1). Comparisons are given for strain *Syn*-sgRNA$_{rbc}$ *cbbM*(WT) at different gas feeds (compare Fig. 2, Supp. Fig. S4). Log2 fold changes (Log2FC) above 1.0 or below -1.0 with associated significant adjusted p values (adj. p < 0.01) were marked in green and red, respectively.

| | | | 5% $CO_2$, 95% $N_2$, 0% $O_2$ / 5% $CO_2$, 75% $N_2$, 20% $O_2$ | | 1% $CO_2$, 78% $N_2$, 21% $O_2$ / 5% $CO_2$, 75% $N_2$, 20% $O_2$ | |
|---|---|---|---|---|---|---|
| | Locus tag | Gene name | Log2FC | adj. p | Log2FC | adj. p |
| Rubisco | *slr0009* | *rbcL* | -0.28 | 4.8E-1 | -0.71 | 1.2E-1 |
| Rubisco | *slr0012* | *rbcS* | -0.36 | 4.7E-1 | -0.46 | 3.8E-1 |
| Carboxysome | *sll1028* | *ccmK2* | 0.34 | 3.9E-1 | 1.25 | 2.6E-2 |
| Carboxysome | *sll1029* | *ccmK1* | 1.03 | 2.4E-1 | 2.41 | 3.4E-2 |
| Carboxysome | *sll1030* | *ccmL* | -0.42 | 2.7E-1 | 0.42 | 3.1E-1 |
| Carboxysome | *sll1031* | *ccmM* | 0.59 | 2.3E-1 | 0.07 | 9.1E-1 |
| Carboxysome | *sll1032* | *ccmN* | 0.01 | 9.8E-1 | 0.16 | 8.4E-1 |
| Carboxysome | *slr0169* | *ccmP* | 0.11 | 9.0E-1 | -0.33 | 7.1E-1 |
| Carboxysome | *slr0436* | *ccmO* | 0.19 | 7.2E-1 | 0.33 | 5.5E-1 |
| Carboxysome | *slr1838* | *ccmK3* | 0.27 | 6.0E-1 | -0.61 | 2.7E-1 |
| Carboxysome | *slr1839* | *ccmK4* | 0.57 | 3.3E-2 | **-1.03** | 9.2E-3 |
| Carbonic anhydrase | *slr1347* | *ccaA* | 1.07 | 1.3E-1 | 0.98 | 2.0E-1 |
| CmpR regulon | *slr0040* | *cmpA* | 0.92 | 5.4E-1 | -1.15 | 4.7E-1 |
| CmpR regulon | *slr0042* | *slr0042* | 0.47 | 6.2E-1 | -1.60 | 1.3E-1 |
| CmpR regulon | *slr0043* | *cmpC* | 0.21 | 5.8E-1 | -1.45 | 1.7E-2 |
| NdhR operon | *sll1594* | *ndhR* | -0.09 | 8.4E-1 | 1.13 | 3.2E-2 |
| NdhR operon | *sll1733* | *ndhD4* | -1.03 | 2.1E-1 | -0.82 | 3.5E-1 |
| NdhR operon | *sll1734* | *cupA* | -0.59 | 4.5E-1 | 0.29 | 7.3E-1 |

| | | | | | | |
|---|---|---|---|---|---|---|
| NdhR operon | *sll1735* | *sll1735* | -0.41 | 5.0E-1 | 0.84 | 2.0E-1 |
| NdhR operon | *slr1512* | *sbtA* | 0.43 | 3.8E-1 | **2.92** | 4.2E-3 |
| NdhR operon | *slr1513* | *sbtB* | -1.16 | 5.5E-2 | 2.07 | 1.6E-2 |
| NdhR operon | *sll0529* | *sll0529* | 0.27 | 2.7E-1 | 1.12 | 1.1E-2 |

**Supp. Table S10:** Proteins with |Log2FC| > 1 and p.adj < 0.01 in comparison of gas feeds of 5% $CO_2$, 95% $N_2$, 0% $O_2$ and 5% $CO_2$, 75% $N_2$, 20% $O_2$ for *Syn*-sgRNA$_{rbc}$ *cbbM*(WT). Subset of Supp. Table S3 showing only the significantly changed proteins.

| Locus tag | Gene name | Gene product | Log2FC | adj. p |
|---|---|---|---|---|
| *slr0935* | *slr0935* | hypothetical protein | 2.2 | 0.007 |
| *sll0565* | *sll0565* | hypothetical protein | 2.1 | 0.007 |
| *slr1261* | *slr1261* | hypothetical protein | 1.7 | 0.008 |
| *sll0002* | *ponA* | penicillin-binding protein, PBP1 | 1.7 | 0.004 |
| *sll1925* | *sll1925* | hypothetical protein | 1.7 | 0.007 |
| *sll0813* | *ctaC* | cytochrome *c* oxidase subunit II | 1.5 | 0.007 |
| *slr1835* | *psaB* | P700 apoprotein subunit Ib | 1.5 | 0.007 |
| *sll1682* | *sll1682* | alanine dehydrogenase | 1.4 | 0.007 |
| *sll0226* | *ycf4* | photosystem I assembly related protein | 1.4 | 0.008 |
| *sll1019* | *gloB* | hydroxyacylglutathione hydrolase | 1.4 | 0.007 |
| *slr1470* | *slr1470* | PSII auxiliary membrane protein | 1.4 | 0.008 |
| *sll1317* | *petA* | apocytochrome *f*; component of cytochrome $b_6f$ complex | 1.3 | 0.007 |
| *sll1316* | *petC2* | cytochrome $b_6f$ complex iron-sulfur subunit 2 | 1.3 | 0.007 |
| *sll1656* | *sll1656* | hypothetical protein | 1.3 | 0.004 |
| *slr0369* | *slr0369* | RND multidrug efflux transporter | 1.3 | 0.008 |
| *sll0672* | *pacL* | cation-transporting p-type ATPase PacL | 1.3 | 0.003 |
| *sll1453* | *nrtD* | nitrate/nitrite transport system ATP-binding protein | 1.3 | 0.007 |
| *slr1790* | *slr1790* | hypothetical protein | 1.2 | 0.007 |
| *slr0906* | *psbB* | photosystem II core light harvesting protein | 1.2 | 0.007 |
| *slr1755* | *gpsA* | NAD+ dependent glycerol-3-phosphate dehydrogenase | 1.2 | 0.007 |
| *slr0657* | *lysC* | aspartate kinase | 1.2 | 0.008 |
| *slr1471* | *yidC* | Membrane protein insertase YidC | 1.2 | 0.001 |

| | | | | |
|---|---|---|---|---|
| *sll0851* | *psbC* | photosystem II CP43 protein | 1.1 | 0.008 |

*Supplementary Tables S11 to S14 are part of the Excel workbook SuppTables_Synechocystis_CbbM-screening.xlsx*

**Supp. Table S15:** EVmutation epistatic predictions for all 16 variants included in the mutant pool used for screening. WT: wild-type CbbM, 1: S162P, 2: A230E, 3: V323A, 4: G422R. Variants are sorted according to the EVmutation epistatic score. This score is calculated based on the residue conservation at a specific site and its couplings to other residue positions.

| Mutant name | Amino acid exchanges | EVmutation epistatic score |
|---|---|---|
| WT | none | 0.0 |
| 3 | V323A | 2.8 |
| 2 | A230E | 2.9 |
| 1 | S162P | 3.1 |
| 4 | G422R | 4.7 |
| 23 | A230E, V323A | 5.8 |
| 13 | S162P, V323A | 6.1 |
| 12 | S162P, A230E | 6.1 |
| 34 | V323A, G422R | 7.6 |
| 24 | A230E, G422R | 7.6 |
| 14 | S162P, G422R | 7.9 |
| 123 | S162P, A230E, V323A | 9.1 |
| 234 | A230E, V323A, G422R | 10.5 |
| 134 | S162P, V323A, G422R | 10.8 |
| 124 | S162P, A230E, G422R | 10.9 |
| 1234 | S162P, A230E, V323A, G422R | 13.9 |

*Supplementary Tables S16 is part of the Excel workbook*

*SuppTables_Synechocystis_CbbM-screening.xlsx*

**Supp. Table S17:** First two inflection points of different CbbM variants as determined by nanoDSF. The mean and standard deviation of three technical replicates is given.

| CbbM variant code | Amino acid exchanges | Inflection point #1 (°C) | Inflection point #2 (°C) |
|---|---|---|---|
| 0 | - | 44.92 ± 0.04 | 70.12 ± 0.06 |
| 2 | A230E | 44.17 ± 0.08 | 69.76 ± 0.03 |
| 4 | G422R | 49.25 ± 0.02 | 70.00 ± 0.06 |
| 14 | S162P, G422R | 49.72 ± 0.14 | 69.27 ± 0.26 |
| 1234 | S162P, A230E, V323A, G422R | 52.63 ± 0.14 | 68.56 ± 0.08 |

**Supp. Table S18:** Oligonucleotides used in this study.

| Name | Sequence | Function |
|---|---|---|
| P01 | GGCCACCGGTGTTGTATTGT | Test of integration of complete CRISPRi construct into genome |
| P02 | CTAGCTCACTCGGTCGCTACTAC | Test of integration of complete CRISPRi construct into genome |
| P03 | CCTGTGGTCACGGTTCTGTT | Test of integration of complete CRISPRi construct into genome |
| P04 | GCTTGCAGCACCAACATGAA | Test of integration of complete CRISPRi construct into genome |
| P05 | TGGGATTGTAACAATTTTTAGTGGTCA | Test of full segregation of *ndhD3* deletion strain |
| P06 | CATGGAGGCATAACCCCGTT | Test of full segregation of *ndhD3* deletion strain |
| P07 | TGCCTACCTGAATCAAACTGTCA | Test of full segregation of *ndhD4* deletion strain |
| P08 | TTTCTTTGGCCGTCTCACCA | Test of full segregation of *ndhD4* deletion strain |
| M01 | GGTTCCTGAGGAAATGGTACGCAAG | Site-directed mutagenesis of *Gallionella* Rubisco coding sequence, M140E |
| M02 | AAGAAGTCCAACATGCGTAAAC | Site-directed mutagenesis of *Gallionella* Rubisco coding sequence, M140E |
| M03 | TTTAGGTCGGCCCGAGACGGACG | Site-directed mutagenesis of *Gallionella* Rubisco coding sequence, S162P |
| M04 | ACTTTCCATAGATTGCTGATATTC | Site-directed mutagenesis of *Gallionella* Rubisco coding sequence, S162P |
| M05 | AGCCCAGCAAGAAACCGGCCAAG | Site-directed mutagenesis of *Gallionella* Rubisco coding sequence, A230E |
| M06 | CGATCCATAGCTTCTGCCAC | Site-directed mutagenesis of *Gallionella* Rubisco coding sequence, A230E |
| M07 | TATGAAGCTCGCCCGCCTAATGG | Site-directed mutagenesis of *Gallionella* Rubisco coding sequence, V323A |
| M08 | TAGCATAGAGGGTCCATG | Site-directed mutagenesis of *Gallionella* Rubisco coding sequence, V323A |
| M09 | TATTAGCCTACGGCAGGCGTATGACTG | Site-directed mutagenesis of *Gallionella* Rubisco coding sequence, G422R |

| | | |
|---|---|---|
| M10 | CCTCCAGCTGCTGGAGAA | Site-directed mutagenesis of *Gallionella* Rubisco coding sequence, G422R |
| S1 | ACTGGAGTTCAGACGTGTGCTCTTCCGATCTTGGCCGCGGTAACTAGTAAGC | PAGE-purified, Illumina library preparation |
| S2 | ACACTCTTTCCCTACACGACGCTCTTCCGATCTNGTCTAGAATCGCCGAAAGTAATTCAACTCC | PAGE-purified, Illumina library preparation |
| S3 | ACACTCTTTCCCTACACGACGCTCTTCCGATCTNNGTCTAGAATCGCCGAAAGTAATTCAACTCC | PAGE-purified, Illumina library preparation |
| S4 | ACACTCTTTCCCTACACGACGCTCTTCCGATCTNNNGTCTAGAATCGCCGAAAGTAATTCAACTCC | PAGE-purified, Illumina library preparation |

**Supp. Table S19:** *Synechocystis* strains used in this study.

| Name | Comment | Description |
|---|---|---|
| *Syn*-sgRNA(-) | Wild-type strain transformed with plasmid pMD19T_psbA1_PL22_dCas9_B0015_SpR from (6), which replaces the *slr1181* (*psbA1*) locus by the gene encoding dCas9 | Δ*psbA1* $P_{J23101}$::*tetR* $P_{L22}$::*SPdcas9* |
| *Syn*-sgRNA$_{rbc}$ | As *Syn*-sgRNA(-), but with two sgRNAs complementary to the coding sequence of *rbcL*. The sgRNA sequences were obtained from the two strongest acting sgRNAs targeting *rbcL* in (7) | Δ*psbA1* $P_{J23101}$::*tetR* $P_{L22}$::*SPdcas9* $P_{L22}$::*sgRNA_rbcL_1* $P_{L22}$::*sgRNA_rbcL_2* |
| *Syn*-sgRNA$_{rbc}$ *cbbM*(WT) | As *Syn*-sgRNA$_{rbc}$, with RSF1010-based plasmid harboring gene encoding wild-type CbbM, *cbbM*$^{+}$, under control of trc promoter. | $P_{trc}$::*(N-Strep)-cbbM*$^{+}$ Δ*psbA1* $P_{J23101}$::*tetR* $P_{L22}$::*SPdcas9* $P_{L22}$::*sgRNA_rbcL_1* $P_{L22}$::*sgRNA_rbcL_2* |
| *Syn*-sgRNA$_{rbc}$ *cbbM*(xxxx) | As *Syn*-sgRNA$_{rbc}$ *cbbM*(WT), but with a mutant variant of CbbM encoded instead of wild-type CbbM. xxxx stands for the combination of introduced amino acid exchanges, encoded in the following: 1: S162P, 2: A230E, 3: V323A, 4: G422R | $P_{trc}$::*(N-Strep)-cbbM(xxxx)* Δ*psbA1* $P_{J23101}$::*tetR* $P_{L22}$::*SPdcas9* $P_{L22}$::*sgRNA_rbcL_1* $P_{L22}$::*sgRNA_rbcL_2* |
| *cbbM*-$GFP_{11}$\|$GFP_{1-10}$ | Wild-type CbbM tagged with 11th alpha helix of split-GFP (8) expressed from trc promoter and encoded on RSF1010-based plasmid, remaining part of GFP (1st to 10th alpha helices (8)) expressed under rhamnose-inducible promoter | Δ*slr0168 cmR* $P_{rha}$::*spGFP1-10* Ptrc::*(N-Strep)-cbbM-spGFP11* |
| *cbbM*-$GFP_{11}$\|noGFP | Wild-type CbbM tagged with 11th alpha helix of split-GFP (8) expressed from trc promoter and encoded on RSF1010-based plasmid, control without remaining part of GFP | Δ*slr0168 cmR* $P_{trc}$::*(N-Strep)-cbbM-spGFP11* |
| *Syn*-sgRNA$_{rbc}$ *cbbM*(WT) Δ*ndhD3* Δ*ndhD4* | As *Syn*-sgRNA$_{rbc}$ *cbbM*(WT), with deletions of genes encoding NDH-1 subunits NdhD3 and NdhD4, encoded by *ndhD3* (*sll1733*) and *ndhD4* (*sll0027*). | $P_{trc}$::*(N-Strep)-cbbM*$^{+}$ Δ*psbA1* $P_{J23101}$::*tetR* $P_{L22}$::*SPdcas9* $P_{L22}$::*sgRNA_rbcL_1* $P_{L22}$::*sgRNA_rbcL_2* Δ*ndhD3* Δ*ndhD4* |

**Supp. Table S20:** Molecular weights and extinction coefficients of the Rubisco mutant variants as calculated using Expasy ProtParam.

| Mutant variant | Molecular weight [Da] | Extinction coefficient |
|---|---|---|
| 0 | 53392.59 | 61685 |
| 4 | 53491.72 | 61685 |
| 2 | 53450.62 | 61685 |
| 14 | 53501.76 | 61685 |
| 1234 | 53531.74 | 61685 |

**Supp. Table S21:** Spectrophotometric Rubisco assay components, preparation and storage. Storage conditions for ATP, phosphocreatine, NADH, creatine phosphokinase, glyceraldehyde 3-phosphate dehydrogenase and 3-phosphoglyceric phosphokinase were inspired by (9).

| Component | Assay concentration | Volume to add | Stock | Water or 1 M EPPS | Storage |
|---|---|---|---|---|---|
| $NaHCO_3$ pH 8.4 | 50 mM | 10 µL | 500 mM | Water | RT |
| $MgCl_2$ | 20 mM | 10 µL | 200 mM | Water | RT |
| DTT | 0.5 mM | 5 µL | 10 mM | Water | -20°C |
| ATP | 2 mM | 5 µL | 40 mM | Water | -80°C |
| Phosphocreatine | 10 mM | 5 µL | 200 mM | Water | -80°C |
| NADH | 0.5 mM | 5 µL | 10 mM | EPPS | -80°C |
| Carbonic anhydrase | 0.1 mg/mL | 10 µL | 1 mg/mL | EPPS | -80°C |
| Creatine phosphokinase | 20 U/mL | 10 µL | 200 U/mL | EPPS | -80°C |
| Glyceraldehyde 3-phosphate dehydrogenase | 20 U/mL | 10 µL | 200 U/mL | EPPS | -80°C |
| 3-Phosphoglyceric phosphokinase | 20 U/mL | 10 µL | 200 U/mL | EPPS | -80°C |
| Ribulose-1,5-bisphosphate | 1 mM | 10 µL | 10 mM | Water | -80°C |
| RuBisCO | 500 nM | 10 µL | 5 µM | 20 mM EPPS +20 mM $MgCl_2$ | 4°C |
| Total | | 100 uL | | | |
